# High throughput PRIME editing screens identify functional DNA variants in the human genome

**DOI:** 10.1101/2023.07.12.548736

**Authors:** Xingjie Ren, Han Yang, Jovia L. Nierenberg, Yifan Sun, Jiawen Chen, Cooper Beaman, Thu Pham, Mai Nobuhara, Maya Asami Takagi, Vivek Narayan, Yun Li, Elad Ziv, Yin Shen

## Abstract

Despite tremendous progress in detecting DNA variants associated with human disease, interpreting their functional impact in a high-throughput and base-pair resolution manner remains challenging. Here, we develop a novel pooled prime editing screen method, PRIME, which can be applied to characterize thousands of coding and non-coding variants in a single experiment with high reproducibility. To showcase its applications, we first identified essential nucleotides for a 716 bp *MYC* enhancer via PRIME-mediated saturation mutagenesis. Next, we applied PRIME to functionally characterize 1,304 non-coding variants associated with breast cancer and 3,699 variants from ClinVar. We discovered that 103 non-coding variants and 156 variants of uncertain significance are functional via affecting cell fitness. Collectively, we demonstrate PRIME capable of characterizing genetic variants at base-pair resolution and scale, advancing accurate genome annotation for disease risk prediction, diagnosis, and therapeutic target identification.

## Main

Advances in genome sequencing have led to the identification of hundreds of millions of genetic variants in the human population, with a fraction conferring risk for common illnesses such as diabetes, neurological disorders, and cancers^1^. A major barrier to understanding the genetic underpinnings of these complex diseases is the paucity of functional annotation for disease-associated variants, especially because such variants are predominantly located within non-coding regions. Growing evidence suggests that non-coding risk variants may contribute to disease pathogenesis by disrupting gene regulation. Even protein-coding variants discovered from individuals with disease are frequently classified as Variants of Uncertain Significance (VUS). Therefore, more precise and higher throughput functional characterization methods for elucidating disease-associated variant function at base-pair resolution, and multiplexed across genomic loci, are necessary to realize the potential of personalized medicine.

The development of genome editing technologies has enabled us to perturb and assess DNA sequences in desired regions at a large scale. However, there are still fundamental barriers to utilizing these methods for precision genome annotation. For example, CRISPRa, CRISPRi, CRISPR deletion, and CRISPR indel have been applied in genetic screening strategies for characterizing both genes and *cis*-regulatory regions^2^, but have failed to pinpoint casual variants for diseases. Traditional methods of characterizing DNA variants (SNPs) by knock-in via homologous recombination are inefficient and low throughput. Base editors also have limitations, introducing specific mutations (C→T, A→G, T→C, G→A) with varied target efficiencies^3^. Thus, there is still a significant deficit in methods for effectively characterizing the roles of putative disease-causing variants in human health and diseases. Robust high throughput methods making desired edits at base-pair resolution are urgently needed to achieve a better understanding of the genetic underpinnings of disease.

Prime editing (PE), a versatile and precise genetic engineering method, has been developed to introduce any type of edit, including point mutation, insertion, and deletion^4^. In particular, PE2, employs the *Streptococcus pyogenes* Cas9 (SpCas9) H840A nickase and Moloney murine leukemia virus (M-MLV) reverse transcriptase. The spacer in the prime editing guide RNA (pegRNA) directs the Cas9 nickase and M-MLV complex to the target site, while the RT template sequence provides the desired editing information. Thus, both targeting and editing information can be easily programmed in the same pegRNA to perform single nucleotide substitution, insertion or deletion. PE3, a newer iteration of PE, can further increase editing efficiency by promoting the replacement of non-edited strands using an additional single-guide (sgRNA) for nicking^5^. Prime editors’ capacity for precision genome editing suggests the possibility of high throughput variant-level genome manipulation. Recently, PE screens were used to identify VUS at the *NPC1* locus based on a lysosome functional assay by transfection of pegRNAs and targeted sequencing of this region^6^. Although transient transfection of PE machinery followed by targeted sequencing of the edited locus enables the identification of editing events, its scope is limited to just that locus, and thus, scaling up for massively parallel assessment of multiple loci is not feasible. Besides increased throughput, improved control of transgene copy number, stable expression of PE machinery, and direct loci comparison are also desired.

Here, we enable high throughput pooled screens of thousands of DNA variants in the human genome by lentiviral delivery of PE, namely PRIME. We demonstrate the utility of PRIME for three different applications, including the saturation mutagenesis analysis of a 716 bp enhancer, the functional characterization of 1,304 breast cancer-associated variants, and the evaluation of 3,699 clinical variants’ impact on cell fitness. Our results establish the generalizability of PRIME for precisely characterizing genetic variants in the human genome.

## Optimization of PE efficiency in mammalian cells delivered by lentivirus

To enable PE screens with delivery by lentivirus, we initially installed PE3 by infecting MCF7 cells using three different viruses: 1) virus expressing Cas9 (H840A) nickase (nCas9) and Moloney murine leukemia virus (M-MLV) reverse transcriptase; 2) virus expressing pegRNA; 3) virus expressing nick sgRNA (ngRNA). Unfortunately, this strategy yielded less than 1% PE efficiency with a relatively high indel rate. This is because of the low efficiency of coinfecting three different viruses in the same cell (**Fig. 1a, Supplementary Fig. 1a**).

**Figure 1.**
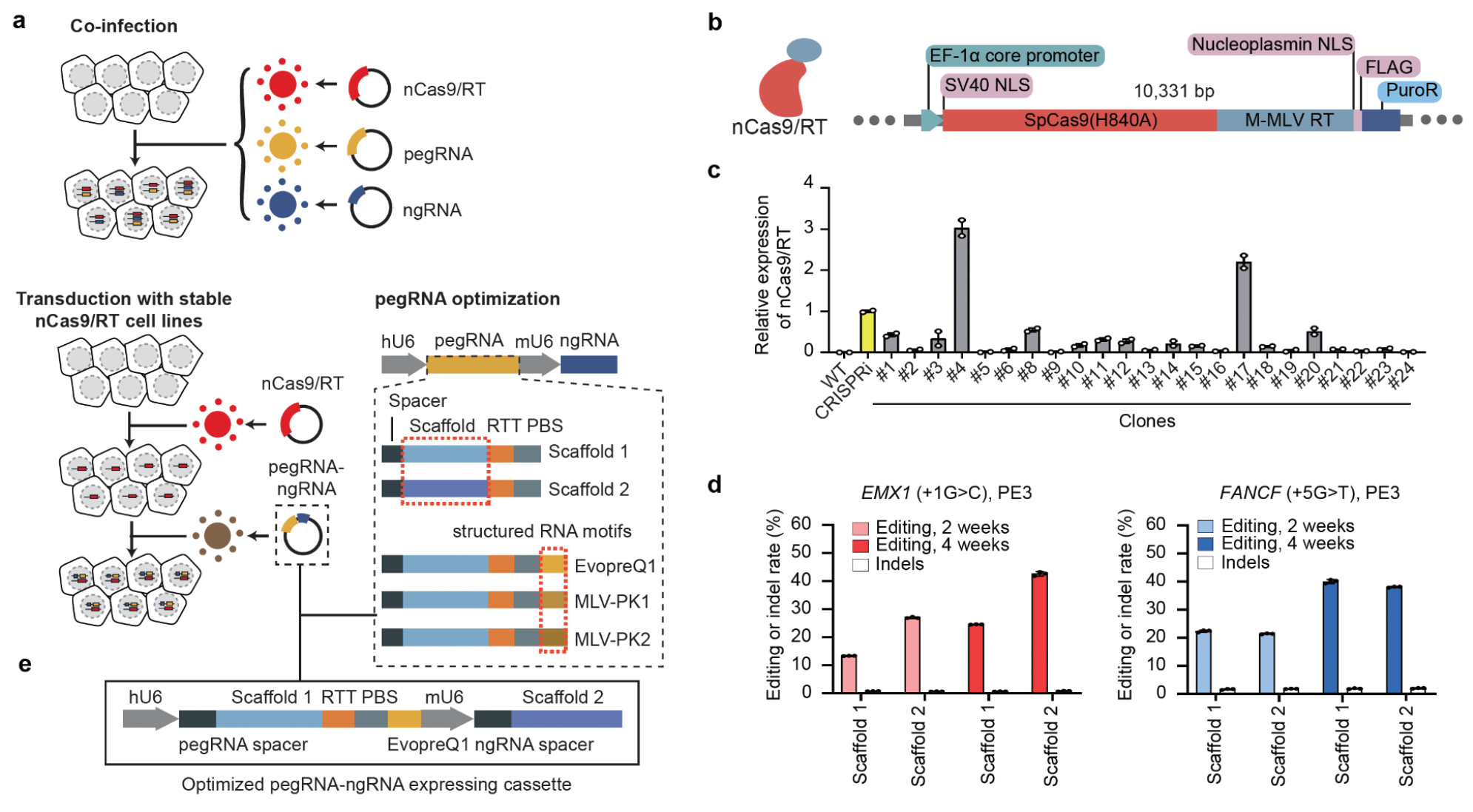
Optimizing PE efficiency in mammalian cells using lentiviral delivery. (a)The different strategies tested for optimizing PE efficiency in MCF7 cell lines. Top: co-infecting three different viruses to deliver PE machinery. Bottom: dual pegRNA/ngRNA viral infection of clonal MCF7 line stably expressing nickase Cas9 (nCas9) and Moloney murine leukemia virus reverse transcriptase (M-MLV RT). Two scaffolds and three different structured RNA motifs tested are also shown. (b) Lentiviral construct for generating nCas9/RT expressing MCF7 clones. PuroR, Puromycin resistance gene. M-MLV RT, Moloney murine leukemia virus reverse transcriptase. (c) RT-qPCR analysis showing the relative expression of nCas9/RT in different clones, normalized to the dCas9 expression of an established CRISPRi iPSC line (Yellow). Error bars represent the s.e.m. (d) The editing efficiency and indel rate for *EMX1* and *FANCF* loci at 2 weeks and 4 weeks after PE installation using two different RNA scaffolds. Error bars represent the s.d. (e) Improved vector for expression of pegRNA and ngRNA for PRIME. RTT: reverse transcription template, PBS: primer binding site.

Packaging all PE3 components within the same virus is challenging. To increase PE efficiency and facilitate a pooled screening approach with a lentiviral library, we infected MCF7 cells with lentivirus containing an nCas9 and M-MLV reverse transcriptase stable expression cassette (**Fig. 1b**). After puromycin selection, we isolated multiple clones and selected one with the highest nCas9 expression (**Fig. 1c**, RT-qPCR, clone #4, **Supplementary Fig. 1b**) for subsequent experiments. The stable expression of nCas9/M-MLV allows for high efficiency pegRNA/ngRNA packaging and lentiviral delivery, with greater editing efficiency than the co-infection method (**Fig. 1d**). To further improve PE efficiency, we assessed editing efficiency using three different structured RNA motifs (EvopreQ1, MLV-PK1, and MLV-PK2) at the 3’ terminus of the pegRNA^7-9^. Cells treated with pegRNAs containing scaffold structure RNA motifs exhibited consistently higher editing efficiencies at both the *EMX1* and *FANCF* locus compared to using PE without structured RNA motifs (**Supplementary Fig. 1c**), so we added evopreQ1 to the pegRNA design for all pooled screens. Scaffold 1^5^ and 2^10^ had no significant effects on PE efficiency, suggesting the feasibility of dual pegRNA and ngRNA delivery from the same viral particle (**Fig. 1d**). All PE experiments in clonal MCF7 cells (MCF7-nCas9/RT) exhibited relatively low indel rates (0.7% to 1.95%). Thus, we used MCF7-nCas9/RT cells and lentiviral delivery of both the pegRNA with scaffold 1 and ngRNA with scaffold 2 in the same construct (**Fig 1e**).

## PRIME enables nucleotide-resolution analyses of enhancer function

Enhancers can modulate cell type-specific gene expression and are highly enriched with disease-associated variants. Knowledge of the endogenous function for each nucleotide in enhancers should reveal crucial transcription factors that govern enhancer activation and facilitate the development of better models for gene regulatory networks and the prediction of disease-associated non-coding variant regulatory effects. To test whether PRIME can quantify the impact of each base in an enhancer, we focused on an MCF7-specific *MYC* enhancer identified from a CRISPRi screen^11^. This enhancer is located 405 kb downstream of *MYC* and displays enhancer signatures, including open chromatin, H3K27ac, and H3K4me1 signals, in addition to forming a chromatin loop with the *MYC* promoter (**Fig. 2a**). Deletion of this enhancer caused an 85% downregulation of *MYC* expression in MCF7 cells confirming its enhancer activity for *MYC* (**Supplementary Fig. 2a**). Since *MYC* downregulation is correlated with MCF7 cell survival^12^, we performed a PE-enabled high throughput saturation mutagenesis screen of this *MYC* enhancer in MCF7 cells dependent on the cell survival phenotype (**Fig. 2b**).

**Figure 2.**
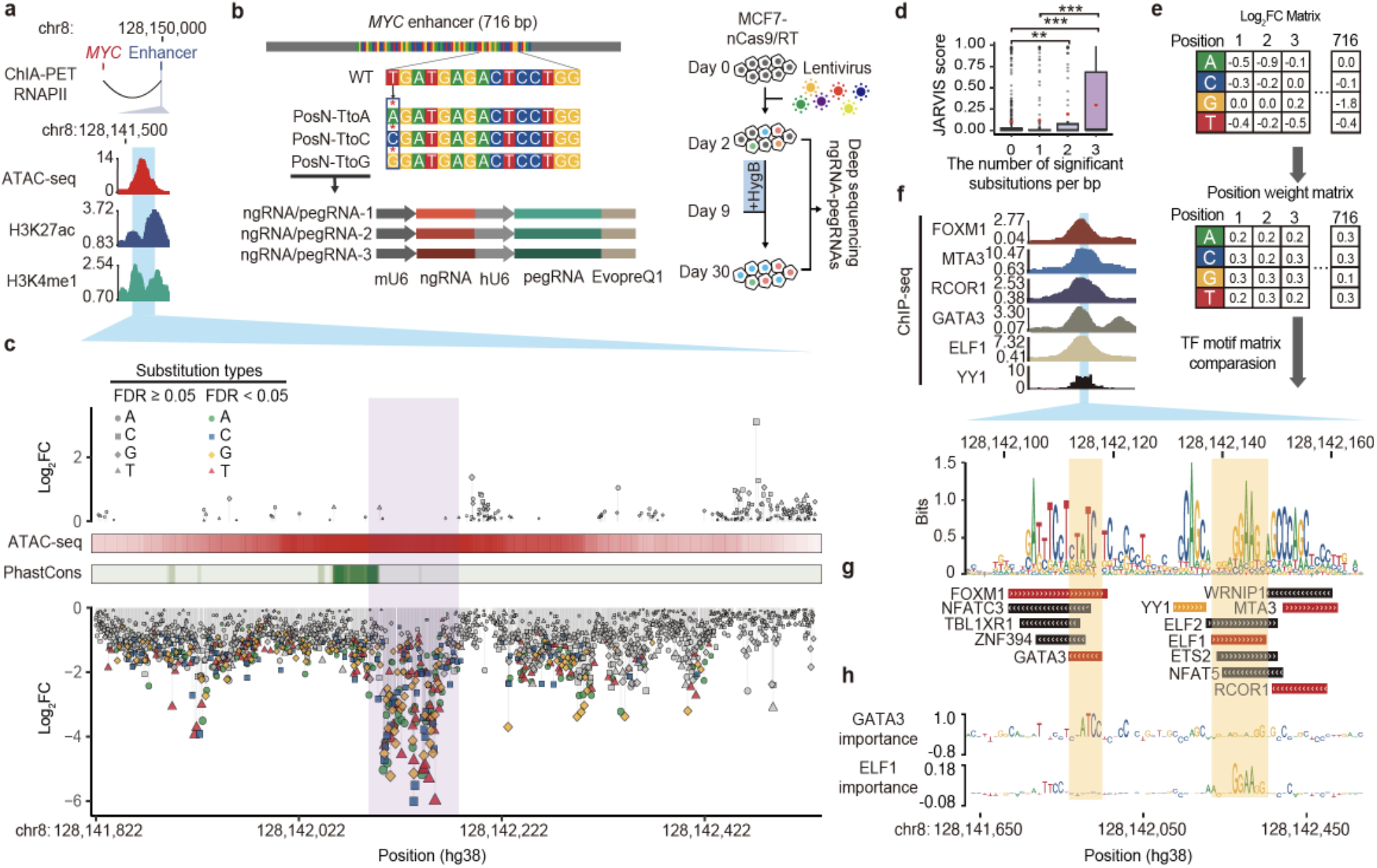
Functional characterization of a *MYC* enhancer by saturation mutagenesis using PRIME. (a) (Top) The target enhancer is downstream of *MYC*. (Bottom) The enhancer region is highly enriched with ATAC-seq, H3K27ac, and H3K4me1 ChIP-seq signals. The blue area indicates the region selected for PRIME. (b) (Top) Diagram showing the design of saturation mutagenesis screening at the 716 bp enhancer. Each nucleotide was subjected to substitution with three nucleotides by PE. (Middle) Each substitution event was covered by three uniquely designed pegRNA/ngRNA pairs. (Bottom) The PRIME workflow. (c) Log2(fold change) of each substitution at each base pair ordered by their genomic locations. Mutations with a significant effect on cell fitness are colored. ATAC-seq signals and conservation scores calculated by PhastCons are shown. (d) JARVIS scores for base pairs with different numbers of significant substitutions. Box plots indicate median, IQR, Q1 − 1.5 × IQR, and Q3 + 1.5 × IQR. Outliers are shown as gray dots. Mean values are shown as red dots. *P* values were calculated using a two-tailed two-sample t-test. (e) The creation of a functional PWM for identifying potential TF binding sites. (f) (Top) ChIP-seq signals of 6 TFs in MCF7. The blue region indicates the core enhancer region. (Bottom) The sequence logo plot for the core enhancer regions generated by the functional PWM from (e). (g) Matched TF binding sites. (h) (Top) Dense tracks showing BPNet model-derived nucleotide importance scores for GATA3 and ELF1 binding sites.

To dissect the enhancer’s function at base-pair resolution, we designed a library of 6,252 pairs of pegRNA/ngRNA to generate 2,148 single nucleotide substitutions within the 716 bp *MYC* enhancer region (**Supplementary Table 1**). Specifically, we changed the original base into three other nucleotides, and each event was independently evaluated three times in the same screen (**Fig. 2b**). We also included 94 positive control pegRNA/ngRNA pairs, which introduced stop codons (iSTOPs) in *MYC*, and 400 negative control pegRNA/ngRNA pairs. 246 of the negative controls were non-human genome targeting, and 154 targeted the *AAVS1* safe harbor locus (**Supplementary Table. 1**). We then infected MCF7-nCas9/RT cells with lentiviral libraries expressing these pegRNA/ngRNA pairs (**Supplementary Fig. 2b**). Two days after infection, virus-transduced cells were hygromycin selected for one week and expanded in regular media for another 3 weeks. We collected cells at 2 and 30 days post-infection, amplified the integrated pegRNA/ngRNA pairs, and determined the relative depletion or enrichment of each pegRNA/ngRNA between these two time points by deep sequencing (**Fig. 2b**). We performed this screen 3 times (**Supplementary Fig. 2c**) and used negative controls, including non-human targeting and AAVS1 targeting paired pegRNA/ngRNAs for data normalization. Fold changes (FC) for each pegRNA/ngRNA pair between day 2 and day 30 samples post-infection were calculated using the MAGeCK pipeline^13^ (**Supplementary Table 1**). As expected, 78% (73/94) of iSTOPs were depleted (log_2_FC < 0) 30 days post-infection. iSTOP depletion rates were negatively correlated with their distance from the transcription start site (TSS) of *MYC*, consistent with the observation that gene knockout is more efficient when perturbations are introduced at the 5’ terminus^14^ (**Supplementary Fig. 2d**). In addition, two iSTOPs (amino acid position 350 and 355) targeting the region between the nuclear localization signal (NLS) and the carboxy-terminal domain (CTD) domain were also significantly depleted (**Supplementary Fig. 2d)**. The N-terminus of MYC contains its core transcription transactivation domain which binds multiple partners^15^. It is possible that those two iSTOPs created a truncated MYC still capable of binding to cofactors, but unable to bind MYC DNA targets, interfering with the functions of wild type MYC and its cofactors.

To investigate the effects of each nucleotide on enhancer function, we defined sensitive base pairs (SBP) as nucleotides that affect cell fitness when substituted at least once (FDR < 0.05, |log_2_FC| > 1). 334 of the 716 (46.6%) tested base pairs were SBP with log_2_FC < -1 (**Supplementary Table 1**), indicating that mutations at those locations reduce enhancer activity and cell fitness. 23.1% (77/334) of SBPs were depleted at day 30 with all three substitutions (FDR < 0.05, log_2_FC < -1). Additionally, none of the tested sequences were significantly enriched at day 30 with increased cell growth phenotype, indicating that perturbation of these sequences exclusively attenuated enhancer activity (**Fig. 2c**).

Deep learning models have been developed to prioritize non-coding regions and predict their relevance to human disease. Encouragingly, SBPs with two or more significant substitutions (n = 172) were predicted to be more deleterious than SBPs with only one significant substitution (n = 162) or non-SBPs (n = 382) by JARVIS^16^ (**Fig. 2d**). This demonstrates the success of PRIME in validating computationally predicted functional sequences. We further established a continuous bin density analysis, detecting variation in SBP density along the enhancer to define SBP-enriched regions (**Supplementary Fig. 2e and f**). We identified the core enhancer region in the enhancer with a high density of SBPs, based on the slope value of the cumulative curve of SBPs with three significant substitutions, as a larger slope value indicates a higher density of SBPs in the region. The core enhancer region was defined by a minimal slope cut-off of 0.43 (Z score-derived *P* < 0.05). The core enhancer region (chr8:128,142,093-128,142,181, hg38) colocalized with an open chromatin summit. This region contains SBPs with the most extensive fold changes when mutated, indicating its strong effect on enhancer activity. (**Fig. 2c**, highlighted in purple). Notably, the enhancer’s core sequence was located next to a highly conserved region (**Fig. 2c**). This is not surprising because enhancers undergo rapid evolutionary changes compared to protein-coding sequences^17^.

Our functional data provide a unique opportunity to calculate and construct a position weight matrix (PWM). Using fold changes for each nucleotide, we generated a functional PWM (**Fig. 2e**). Comparing our functional PWM with curated transcription factors (TFs) motifs from the JASPAR, HOCOMOCO, and SwissRegulon databases^18-20^ identified 13 TFs with matched motif PWMs (**Fig. 2g and h, Supplementary Table 2**). 5 predicted TFs (GATA3, ELF1, FOXM1, MTA3 and RCOR1) have already been shown to bind to the *MYC* enhancer based on ENCODE ChIP-seq datasets^21^, and YY1 is predicted to bind to this enhancer in MCF7 by Avocado through the ENCODE project^22^ (**Fig. 2f**). Furthermore, *GATA3* and *YY1* are essential cell survival genes in MCF7^23^, confirming the utility of PE-enabled saturation mutagenesis for interrogating enhancer function at base pair resolution. Essential nucleotides for the ELF1 and GATA3 binding motifs identified by our screens were consistent with those imputed by BPNet^24^, further validating the importance of quantitative roles of each nucleotide discovered by PRIME. Combined, we demonstrated that PRIME is useful for elucidating nucleotide-resolution functional annotations of non-coding cis-regulatory elements.

## Characterization of breast cancer-associated variants

Next, we tested the feasibility of characterizing >5,000 disease-associated DNA variants at various genomic loci, including non-coding variants from GWAS and variants detected from clinical samples. For GWAS-identified variants, we focused on breast cancer, the most common cancer in women in the U.S. To test the feasibility of characterizing DNA variants associated with breast cancer, we used the summary statistics from the largest GWAS to date, including samples of mostly European ancestry^25^. Candidate genes from a comprehensive fine mapping effort for this GWAS^26^ overlapping with growth phenotype genes prioritized by CRISPR screens^23, 27^ were selected. These include: *CCND1*, *PSMD6*, *MYC*, *UBA52*, *DYNC1I2*, *ESR1*, *MRPS18C*, *NOL7*, *EWSR1*, *BRCA2*, and *GRHL2*, which were negatively selected in a CRISPR knockout screen, and *CUX1*, *CASP8*, and *TNFSF10*, which are tumor suppressor genes and positively selected in a CRISPR knockout screen (**Supplementary Fig. 3a**). We then selected 1,304 single nucleotide polymorphisms (SNPs) (**Supplementary Fig. 3b** and **Supplementary Table 3**) within 500 kbp upstream and downstream of these genes that were previously associated with breast cancer^25^ and had been implicated as possibly acting through these genes^26^. We also selected 3,699 variants from the ClinVar database (**Supplementary Fig. 3c)**, 2,840 of which were identified from patients who were tested for hereditary breast cancer^28^. To systematically assess variants’ impact on cell fitness, we designed two libraries: one to introduce reference alleles (Ref library) and another to introduce alternative alleles (Alt library) targeting the selected variants (**Fig. 3a**) (**Supplementary Table 3**). 250 non-targeting pegRNA/ngRNA pairs were added as negative controls, respectively. For the Alt library, 115 pegRNA/ngRNA pairs introducing stop codons (iSTOPs) in 23 MCF7 growth-related genes were included as positive controls, while pegRNA/ngRNA pairs introducing reference sequences were used for those loci in the Ref library. The cloned plasmids were packaged into lentiviral libraries and transduced into MCF7-nCas9/RT cells. Cells were collected 2 and 32 days post infection, and pegRNA/ngRNA pairs were amplified and deep sequenced (**Fig. 3b**). PRIME replicates using either Ref or Alt library (n = 4) were reproducible at the read count level (**Supplementary Fig. 4a**).

**Figure 3.**
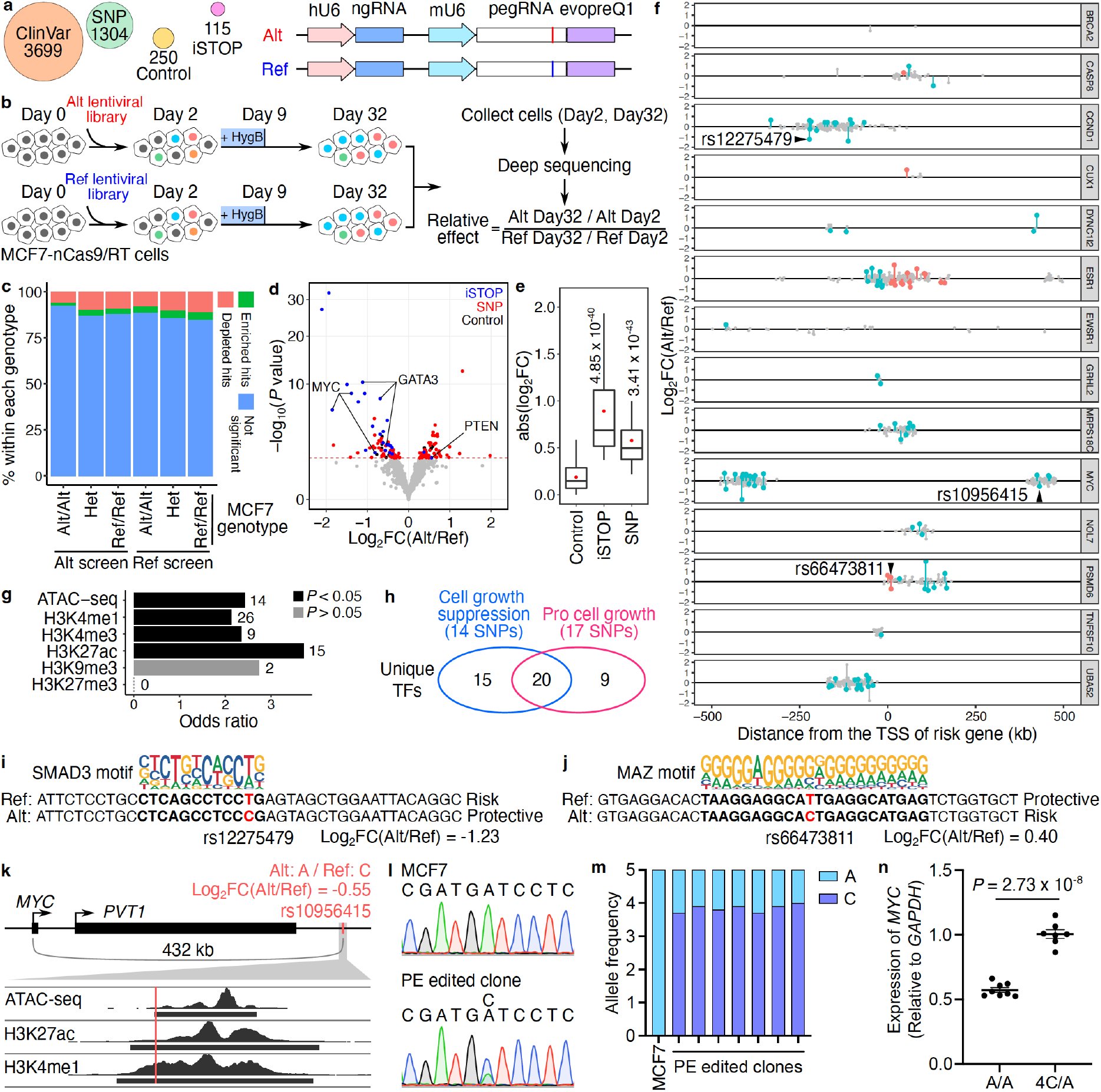
PRIME reveals functional SNPs associated with breast cancer. (a) Alt and Ref library design overview. In the design, we included breast cancer-associated variants (SNP), clinical variants (ClinVar), introduced stop codons (iSTOP), and non-targeting controls. For each variant, pegRNA/ngRNA pairs introducing either the Alt or Ref allele were designed. (b) Workflow of PRIME with Alt and Ref libraries. MCF7-nCas9/RT cells were infected with either lentiviral library. Cells were collected on days 2 and 32 post-infection. The abundance of pegRNA/ngRNA pairs in the samples collected on days 2 and 32 were deep sequenced. The relative effect of each variant was determined based on its relative impact on cell growth between Alt versus Ref alleles. (c) The percentage of significant hits (FDR < 0.05) identified from Alt and Ref screens for Alt/Alt, Het, and Ref/Ref genotypes in MCF7. (d) The functional SNPs (red) with either a positive or a negative impact on cell growth were determined by their relative effect in the Alt versus Ref screens. Blue dots represent significant iSTOPs, and black dots represent controls. The red dashed line indicates 0.05 FDR. (e) Absolute effects of identified functional iSTOPs and SNPs are higher than the effects of negative controls (*P* values were calculated by two-tailed two-sample t-test). (f) The genomic distance of SNPs tested at each risk locus relative to each gene’s TSS. Red dots are functional SNPs within gene bodies, blue dots are functional SNPs in distal regions, and gray dots are SNPs with non-significant effects. (g) Relative enrichment of genomic features for identified functional SNPs (*P* values were calculated by two-tailed Fisher’s exact test). The numbers of SNPs overlapping each genomic feature are labeled next to each bar. (h) Venn diagram showing the numbers of unique transcription factors (TFs) with differential binding sites centered on functional SNPs. The numbers of SNPs that alter TF binding sites are also in the parentheses. (i, j) Examples of functional SNPs disrupting TF binding sites. (i) The Alt protective allele of rs12275749 (position shown in f) affects the SMAD3 binding site and (j) The Alt risk allele of rs66473811 (position shown in f) is matched with the MAZ binding motif. (k) rs10956415 located within a candidate enhancer region overlapping with ATAC-seq, H3K27ac and H3K4me1 peaks in MCF7 cells. (l) Representative Sanger sequencing results for the rs10956415 locus in unedited MCF7 cells and a PE edited clone. (m) Allele frequencies of alternative (A) and reference (C) alleles of rs10956415 in unedited MCF7 cells and PE edited clones. (n) Relative *MYC* expression in control clones and PE edited clones (*P* = 2.73 × 10^-8^, two-tailed two-sample t-test).

From Alt library screens, 33.04% (38/115) of iSTOPs showed a significant cell fitness effect (FDR < 0.05), which is comparable to the 31.8% positivity rate of iSTOPs for common essential genes reported from the base editing screen in MCF7 cells^29^. Furthermore, the fold changes for iSTOPs were highly correlated with those for sgRNAs from MCF7 CRISPR knockout screens of the same genes^23^ (**Supplementary Fig. 4b**). More pegRNA/ngRNA pairs were depleted (FDR < 0.05, Alt screen n = 322 and Ref screen n = 337) than enriched (FDR < 0.05, Alt screen n = 148 and Ref screen n = 209) (binomial test, *P* = 4.78x10^-8^ for Alt screen and *P* = 6.85x10^-16^ for Ref screen) for both Alt and Ref screens on day 32 compared to day 2 (**Supplementary Fig. 4c and d, Supplementary Table 4, 5)**. Theoretically, when a designed peg/ngRNA pair matches the wild type MCF7 genotypes, they should have no effect on cell growth. Notably, however, certain pegRNAs matching the wild type MCF7 genotype, exhibited significant effects on cell growth beyond what was predicted, while the proportion of significant hits for each genotype group were independent of initial MCF7 genotypes (Chi-square test *P* = 0.9998 on the Ref library and *P* = 0.999 on the Alt library, Cochran-Mantel-Haenszel test *P* = 0.9665 for the Ref library and Alt library together). For example, in the Ref library, 11.2% (59 out of 528) of pegRNAs at sites with a Ref/Ref MCF7 genotype exhibited significant depletion, similar to the 10.2% (55 out of 540) at heterozygous sites and 7.9% (18 out of 227) at Alt/Alt genotype sites (**Fig. 3c**). These changes at sites where alleles were not expected to change suggests the presence of undesired consequences of constitutive nCas9 expression, similar to CRISPR inhibition (CRISPRi) once editing machinery is recruited to target sites^30^. To test for potential CRISPRi activity of nCas9 in PE, we compared the results between iSTOPs in the Alt library and the corresponding pegRNA/ngRNA pairs in the Ref library. While pegRNAs in the Ref library exhibited smaller effects on Day 32 compared to iSTOPs targeting the same loci, they were still depleted on Day 32, confirming unintended consequences due to nCas9 occupancy at target genomic loci (**Supplementary Fig. 4e**). Combined, we found that prolonged PE expression exhibits undesired activity similar to CRISPRi, a crucial factor for consideration when analyzing lentivirus-mediated PE screens.

To correct for this undesired PE activity, we compared the ratio of FC for each pegRNA/ngRNA pair from Alt and Ref screens by DESeq2^31^. We determined functional SNPs based on their relative impact on cell growth between Alt and Ref PEs. In total, 56 SNPs with Ref alleles and 47 SNPs with Alt alleles were identified to promote cell growth (*P* < 0.05, empirical significance threshold to control type-I error at 5%, **Supplementary Fig. 4f, Fig. 3d**, and **Supplementary Table 4)**. As expected, identified functional SNPs had smaller effect sizes than stop codons and significantly larger effect sizes than negative control PEs (**Fig. 3e**). Additionally, iSTOPs for genes promoting cell growth, such as *MYC* and *GATA3*, were depleted, while the iSTOP for the cell growth suppressor *PTEN* was enriched, validating our analysis approach (**Fig. 3d**).

Since risk variants can either be the Ref or Alt allele, we further annotated functional SNPs based on genetic annotation of breast cancer risk variants. Since most GWAS SNPs are likely not causal, we expected that only a fraction of the 1,304 tested SNPs would exhibit a biological effect. We calculated the mean likelihood of a variant being causal using CAVIAR and found that the mean expectation for a variant being causal was ∼8.9% when we made the assumption of only one causal variant in each linkage disequilibrium (LD) clump. If we allowed for more than one causal variant in each LD clump the mean probability of being causal for the variants was ∼13.0%. Compared to the reference allele, 50 risk SNPs’ alternative alleles were pro-growth, and 53 risk SNPs’ alternative alleles reduced cell growth (**Fig. 3f**). 18.45% (19/103) of the functionally validated risk SNPs were located within the risk gene’s body. The rest were located in distal regions with an average distance of 185.8 kb from the risk gene’s TSS (**Fig. 3f**). All tested loci contained at least one SNP with a significant effect on cell growth, except for the *BRCA2* locus, in which only 2 SNPs were tested. Finally, identified functional SNPs were significantly enriched for active chromatin marks (two-tailed Fisher’s exact test, *P* < 0.05), including ATAC-seq, H3K27ac, H3K4me1, and H3K4me3 signals, relative to their corresponding genomic background (1 Mbp surrounding selected cell growth genes) (**Fig. 3g**).

To explore potential mechanisms for functional SNPs’ regulation of cell fitness changes, we searched candidate TF binding motifs against the human motif database HOCOMOCO^19^ using 40 bp regions centered on 103 identified functional SNPs. We retrieved 281 and 391 motifs (FDR < 0.05 and TF expression > 1 FPKM) containing Alt and Ref alleles, respectively. After removing redundant motifs for each SNP locus, we identified 90 TF binding sites for 35 unique TFs associated with the cell growth suppression phenotype (log_2_FC(Alt/Ref) < 0) and 55 sites for 29 unique TFs associated with the pro cell growth phenotype (log_2_FC(Alt/Ref) > 0) (**Fig. 3h** and **Supplementary Table 6**). In particular, the Alt allele (protective allele), rs12275479 (T>C) at the *CCND1* locus disrupts the SMAD3 binding motif and is associated with reduced cell growth in our screens, consistent with the TGFβ-SMAD3 axis decreased the number of mammosphere-initiating cells in MCF7^32^ (**Fig. 3f** and **i**). In another example, we found that a MAZ binding site of MAZ is affected by the rs66473811 (T>C) Alt allele at the PSMD6 locus. MAZ is a transcription factor that promotes breast cancer cell proliferation via driving tumor-specific expression of *PPARγ1* gene and regulating *MYC* expression^33, 34^ in line with that Alt allele being the risk allele (**Fig. 3f** and **j**). To validate our PRIME results, we selected rs10956415 from the *MYC* locus, which exhibited a moderate effect on cell growth in the screen (Log_2_FC(Alt/Ref) = -0.55) (**Fig. 3f**). rs10956415 is located in a candidate enhancer region 432 kb downstream of *MYC* (**Fig. 3k**). MCF7 cells are homozygous for the alternative allele (A) at the rs10956415 locus, which has a copy number of five in this cell line^35^ (**Fig. 3l** and **m**). Using prime editing, we converted 4 copies of the alternative allele (A) to the reference allele (C) in seven independent clones, yielding a 43.2% average increase in *MYC* expression compared to unedited cells with 5 copies of A alleles (**Fig. 3m** and **n**). Since *MYC* expression level positively correlates with MCF7 cell growth^36^, the *MYC* expression of the PE edited clones is consistent with the cell growth phenotype of rs10956415 observed in the screening. Together, these results support the use of PRIME to functionally characterize GWAS-identified variants.

## PRIME can characterize clinical variants of uncertain significance

Genetic variants detected in clinical samples provide a valuable resource for understanding the etiologies of human diseases. However, many clinically discovered variants are annotated as Variants of Uncertain Significance (VUS) due to unpredictable functional consequences, even in well-characterized protein-coding genes. To assess the capacity of PRIME to functionally annotate VUS using MCF7 growth phenotypes, we designed pegRNA/ngRNA pairs for 2,532 VUS, 745 pathogenic variants, and 422 benign variants for 17 genes (**Supplementary Fig. 3c** and **Supplementary Table 3)**. 76.78% of the variants tested were from breast cancer patients (**Supplementary Table 3**). By comparing the relative effect sizes of each Alt and Ref allele pair, we identified 236 functional clinical variants affecting cell growth in 15 genes, including 49 pathogenic variants, 156 VUS, and 31 benign variants (**Fig. 4a** and **Supplementary Table 5**). The average effect sizes for pathogenic variants, VUS, and benign variants were between that of negative controls and iSTOPs (**Fig. 4b**).

**Fig 4.**
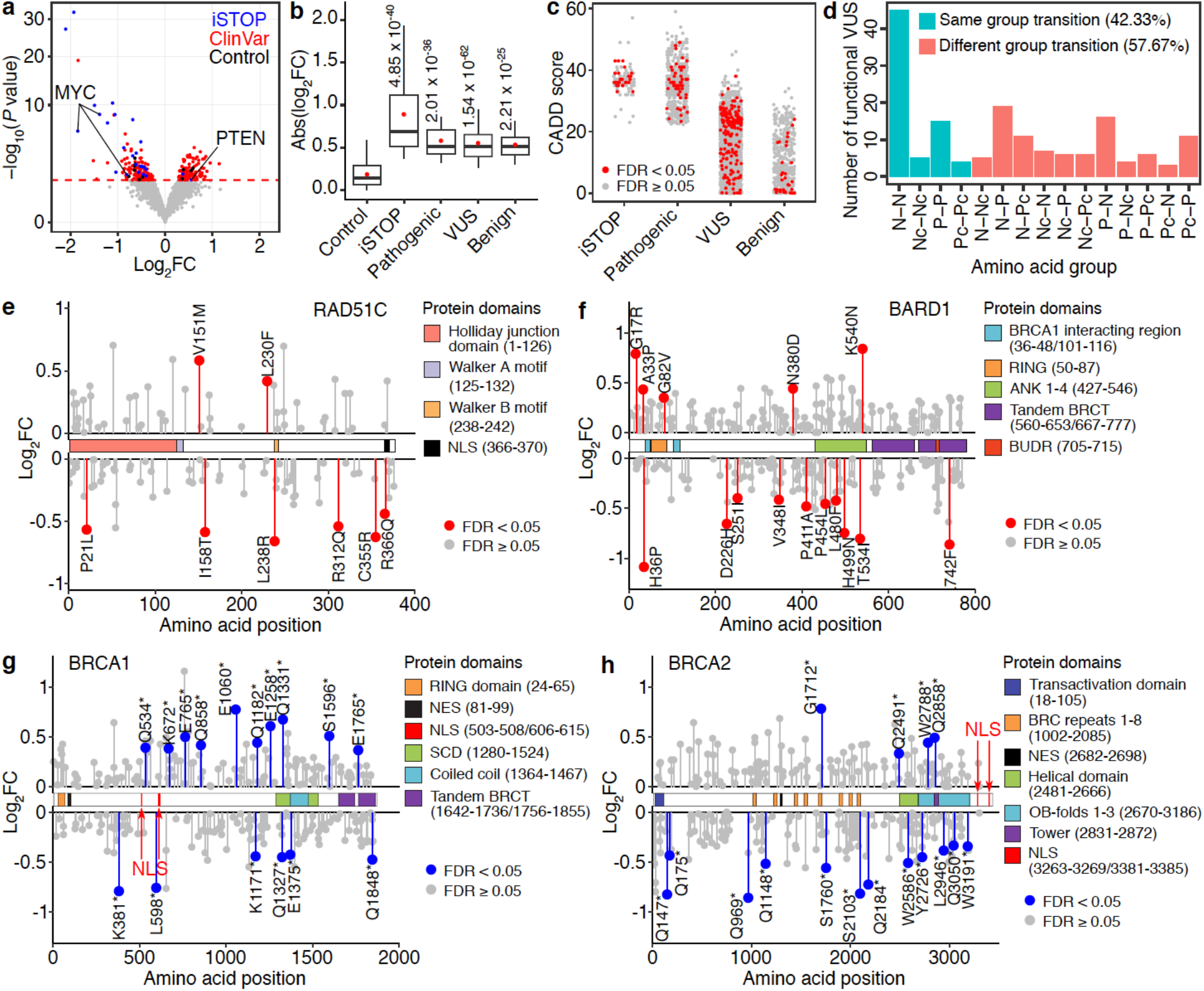
Functional clinical variants identified using PRIME. (a) Functional clinical variants (red) with either a positive or a negative impact on cell growth were determined by relative effects on cell fitness between Alt and Ref alleles. Blue dots represent significant iSTOPs, and black dots represent negative controls. The red dashed line indicates 5% FDR. (b) Effect sizes of identified functional iSTOPs and clinical variants are larger than that of negative controls (*P* values were calculated by two-tailed two-sample t-test). Box plots indicate the median, IQR, Q1 − 1.5 × IQR, and Q3 + 1.5 × IQR. Red dots indicate the mean. (c) CADD scores for iSTOPs and clinical variants. (d) Number of identified functional VUS causing each amino acid group transition. (N, Nonpolar; P, Polar; Pc, Positively charged; Nc, Negatively charged). (e,f) Lollipop plots of functional VUS in *RAD51C* and *BARD1* mapped to their canonical isoforms. The identified significant VUSs are labeled in red. Their effects on cell growth are indicated by fold changes. (g,h) Lollipop plots of the nonsense variants in *BRCA1* and *BRCA2* mapped to their canonical isoforms. The identified significant hits are labeled in blue. Their effects on cell growth are indicated by fold changes.

Several computational metrics have been used to assess the deleteriousness of variants^37, 38^. One such method is CADD, which integrates diverse genome annotations into a single, quantitative score estimating the relative pathogenicity of human genetic variants^37^. iSTOPs and pathogenic variants have similarly high CADD scores relative to other categories (**Fig. 4c**). The CADD scores for the VUS and benign variants exhibit a broad distribution with median scores much lower than those of iSTOPs and pathogenic variants. Interestingly, the CADD scores for identified functional variants within the VUS or benign variant groups did not have higher CADD scores as expected, indicating the limitation of solely relying on computational prediction for variants annotation and underscoring the importance of validating clinical variants with functional assays, even for those located in well-studied protein-coding genes. For example, one benign variant in BARD1 (Arg378Ser) with a low CADD score (CADD = 4.317) would not be classified as functional. However, this variant exhibited a significant cell growth suppression effect in MCF7 cells based on our screening results. BARD1 (Arg378Ser) can impair the nuclear localization of the BRCA1/BARD1 complex, and synergistically promote tumor formation with BARD1 (Pro24Ser) *in vivo*^39^. Furthermore, most of the identified functional VUS were missense variants, and about half of the significant VUS from our screens changed amino acid type within the same group based on polarity (**Fig 4d**), complicating the determination of their molecular consequences. Our results offer novel insights into the potential roles of clinical variants in disease pathogenesis through their modulation of cell fitness, and provide annotations for VUS and benign variants previously uncharacterized.

Functional and structural domains are integral contributors to protein function. 60% of the functional VUS identified are located within an annotated protein domain in the UniProt database^40^, supporting their pathogenicity. For example, we identified 8 VUS in *RAD51C* (**Fig. 4e**), a cancer susceptibility gene and an essential gene for MCF7 survival. Two variants, one (Pro21Leu) in the RAD51C functional domain (amino acid: 1-126) for Holliday junction processing and the other (Arg366Gln) in the NLS region (amino acid: 366-370), were associated with reduced cell growth by our screens (**Fig. 4e**). We also identified functional variants that were not located in any annotated domain, including a functional RAD51C VUS (Arg312Gln) associated with a phenotype of reduced MCF7 growth (**Fig. 4e**). Since Arg312Trp in RAD51C results in homologous recombination deficiency and reduced colony formation phenotypes in MCF10A cells, and abolishes RAD51C-RAD51D interaction^41^, Arg312Gln may produce a similar pathogenic consequence on protein function. When comparing the RAD51C sequence with other RAD51 family proteins, we observed functional VUS were located in both conserved and non-conserved amino acids (**Supplementary Fig. 5a**), underscoring the challenge of predicting variant function based solely on protein sequence conservation.

Protein-protein interaction (PPI) is another essential functional activity in many biological processes. In this study, we also identified functional VUS located in protein binding regions with the potential to affect PPI. For example, BARD1 interacts with BRCA1 through RING domains, and BRCA1-BARD1’s ubiquitin ligase activity is indispensable for DNA double-strand break repair^42, 43^. We identified a functional VUS (His36Pro) in the BARD1 RING domain (**Fig. 4f**), suggesting the structural consequences of this clinical variant affecting BARD1-BRCA1 heterodimer formation (**Supplementary Fig. 5b**). Consistent with these findings, AlphaFold predicts that the His36Pro variant disrupts hydrogen bond formation between His36 in BARD1 and Asp96 in BRCA1 (**Supplementary Fig. 5c**).

Nonsense mutations can generate new stop codons and truncated proteins. Although most are annotated as pathogenic variants in ClinVar, the functional consequences of many remain uncharacterized^28^. In our screens, 563 nonsense clinical variants were tested in 13 breast cancer risk genes with 38 variants identified as positive hits in 7 genes. Remarkably, 39.47% (15/38) exhibited unexpected phenotypes compared to the knockout phenotypes of cell death of these genes. Specifically, a similar number of functional nonsense variants in *BRCA1* (n = 15) and *BRCA2* (n = 16) (**Fig. 4g, h**) were identified; however, 60% (9/15) in *BRCA1* could promote MCF7 cell growth compared to 25% (4/16) in *BRCA2*. After locating variants within BRCA1 and BRCA2, we noticed that truncated proteins resulting from all gain-of-function nonsense variants in BRCA1 still retained their NLS. These results were confirmed by a different nonsense mutation at Q858, located downstream of the NLS in BRCA1, which resulted in truncated BRCA1 with NLS and increased cell growth of MCF7^29^. However, for all of the functional variants identified in BRCA2, their NLSs were located at the c-terminus^44^ and were thus removed from the truncated proteins, leading to the loss of BRCA2 nuclear localization. Collectively, these results demonstrate the capability of PRIME to functionally characterize some nonsense mutations.

## Discussion

In this study, we describe a new genomic screening method, PRIME, to interrogate DNA function at base-pair resolution by adopting and optimizing ‘search-and-replace’ prime editing^5, 9^. We demonstrate the success of pooled prime-editing screens to identify essential nucleotides in a *MYC* enhancer via saturation mutagenesis screen, the functional characterization of 1,304 breast cancer-associated risk SNPs, and provide accurate annotation for 3,699 clinical variants. Our study offers a novel strategy to elucidate genome function at an unprecedented precision and scale. The broad applications demonstrated in this work suggest that PRIME can significantly augment the functional characterization toolbox and advance our ability to elucidate the roles of disease-associated variants in the human genome.

Our analyses show that lentiviral installation of PE yields long-lasting expression of nCas9, pegRNA, and ngRNAs, but can result in unwanted sequence-specific repression similar to CRISPRi. This bias must be corrected to produce accurate base-pair resolution annotations. When assessing the functional impact of a variant, pegRNA controls should be included to introduce other alleles at the same locus. Our study normalized sequence-specific repression bias by comparing the differential effects on cell survival of all base pair substitutions at each locus in the *MYC* enhancer, and between Alt and Ref alleles for disease variants. Additional improvement could be achieved through controlled nCas9 expression duration. For example, a doxycycline-inducible nCas9 could be selectively expressed when editing is needed and reversibly turned off afterwards. In addition to establishing and optimizing PRIME, we defined sensitive base pairs (SBPs) and core sequences for a *MYC* enhancer’s function. We generated a functional PWM for this enhancer by leveraging effect sizes for all possible substitutions at each base from the screens. The functional PWM enabled us to accurately predict TF binding sites within the enhancer, providing critical annotations for delineating *MYC* activation in MCF7 cells. Interpreting the effect of inherited genetic variations will dramatically advance our ability to predict an individual’s disease risk. However, utilizing GWAS data for risk prediction is still limited without substantial functional annotation. In this study, 7.9% of the 1,304 tested GWAS breast cancer variants, and 6.2% of the 2,532 tested VUS were identified as significant hits with functions linked to MCF7 growth phenotypes. Our results demonstrate the feasibility of PRIME for functionally characterizing individual variants. The impact of variants was context-specific and our findings were limited to assessing variants with growth phenotype related functions in MCF7 cells. Other ClinVar did not show changes in our functional assay likely have functional consequences for breast cancer susceptibility genes in a different cell type or other biological processes.

Future work employing different phenotypic screening readouts across multiple cell lines will provide new insights into variant function. For example, screens that identify variants associated with differential drug treatment responses will help construct better predictive models for an individual’s unique benefits and risks from therapeutics. Screens of variants with readouts directly linked to physiological functions e.g. endolysosomal activities in microglia or synaptic activities in neurons using iPSC models will uncover functional variants associated with neuropsychiatric diseases. In summary, our study provides a roadmap to advance functional genomics toward the actionable disease prediction, prevention and treatment necessary to realize personalized medicine.

## Acknowledgements

We thank Howard Y. Chang for sharing wild type MCF7 cells. This work was supported by the Laboratory for Genomic Research Innovation Award LGRFU1019 and the George and Judy Marcus Innovation Fund - Precision Medicine Innovation SBI2019 (to Y.S. and E.Z), the National Institutes of Health (NIH) grants R01AG057497, R01EY027789, UM1HG009402, and U01DA052713 (to Y.S.), U01HG011720 (to Y.L.). Sequencing was performed at the UCSF CAT, supported by UCSF PBBR, RRP IMIA, and NIH 1S10OD028511-01 grants.

## Author contributions

X.R. H.Y., and Y.S. conceived the study. Y.S. and E.Z. supervised the study. X.R. and H.Y. designed PRIME screens. X.R. H.Y. C.B. Y.S. M.N. M.A.T. and V.N. performed experiments under the supervision of Y.S. X.R. HY, J.L.N, Y.S. and J.C. performed computational analysis under the supervision of Y.S. Y.L. and E.Z. Y.S. X.R. and H.Y. prepared the manuscript with input from all other authors.

## Code availability statement

A copy of the custom code used for data analysis and figure generation in this study is available upon request.

## Supplementary Tables

**Supplementary Table 1. pegRNA and ngRNA oligo sequences and their fold changes in *MYC* enhancer.**

**Supplementary Table 2. TF motif analysis for alleles based on functional data from PRIME. Supplementary Table 3. pegRNA and ngRNA oligo sequences for SNP and ClinVar.**

**Supplementary Table 4. PRIME results for breast cancer-associated variants. Supplementary Table 5. PRIME results for clinical variants.**

**Supplementary Table 6. TF motif analysis for alleles with functional SNPs for breast cancer.**

## Methods

### Cell culture

MCF7 cells were cultured in Dulbecco’s Modified Eagle Medium (DMEM) (Gibco, 10569010) supplemented with 10% fetal bovine serum (FBS) (HyClone, SH30396.03), and were passaged with trypsin-EDTA (Gibco, 25200072). All cells were cultured with 5% CO_2_ at 37°C and verified to be free of mycoplasma using the MycoAlert Mycoplasma Detection Kit (Lonza, LT07-218). Wild type MCF7 cells were a gift from Howard Y. Chang’s lab. The MCF7-nCas9/RT cell line was generated by lentiviral transduction of cells with a cassette expressing the nickase Cas9 (nCas9) Moloney murine leukemia virus reverse transcriptase (M-MLV RT) fusion protein. The infected MCF7 cell pool was treated with puromycin (2.5 µg/ml) for two weeks. Then, single cells were sorted into 96-well plates with one cell per well by fluorescence-activated cell sorting (FACS) to generate a clonal MCF7-nCas9/RT cell line. nCas9/RT expression levels were quantified in each clone via RT-qPCR, and normalized to the dCas9 expression level in a WTC11 doxycycline-inducible dCas9-KRAB iPSC line^45, 46^.

### Functional characterization of a *MYC* enhancer by CRISPR deletion

Two sgRNAs were designed to knock out a MCF7 enhancer (chr8:128,141,747-128,142,627, hg38) (sg1: GAAGTTGTAAGTATAGCGAG, sg2: AGTGCCTGGCACAAGGCAGA). SgRNAs were synthesized *in vitro* using the Precision gRNA Synthesis Kit (Invitrogen, A29377) according to the manufacturer protocol and concentrations were quantified with Nanodrop. To deliver genome editing machinery, 100 pmol of Cas9-NLS protein (QB3 MacroLab in University of California, Berkeley) and 120 pmol of *in vitro* synthesized gRNA were electroporated into 250,000 MCF7 cells with the P3 primary nucleofection solution (Lonza, V4XP-3024), using the DN-100 Lonza 4D-Nucleofector program. Cells were then plated into 6-well plates and cultured for 2 days, followed by plating into 96-well plates to pick single clones. Successful knockout clones were identified by genomic PCR with the primers forward: CACCAGGACTTGAAGGCAGC and reverse: CACTTCCCAACCTCAGTTTCC. RT-qPCR was used to quantify *MYC* expression (*MYC* forward primer: GTCCTCGGATTCTCTGCTCT, reverse primer ATCTTCTTGTTCCTCCTCAGAGTC) and normalized to the *GAPDH* expression level (*GAPDH* forward primer: ATTCCATGGCACCGTCAAGG, reverse primer TTCTCCATGGTGGTGAAGACG).

### Cloning of prime editing plasmids

To construct the lentiV2-EF1a-nCas9/RT plasmid, we first excised the U6-sgRNA cassette from the lentiCRISPR v2 plasmid (Addgene, 52961) by dual KpnI and EcoRI digestion followed by blunt end ligation. We further replaced the Cas9 cassette with an nCas9/M-MLV-RT cassette from the pCMV-PE2 plasmid (Addgene, 132775). The lentiV2-pegRNA and lentiV2-ngRNA plasmids were constructed by replacing the Cas9 and Puromycin sequences in the lentiCRISPR v2 plasmid (Addgene, 52961), with hygromycin B and EGFP sequences. RNA motifs and sgRNA scaffolds were further integrated by Gibson assembly.

### Testing prime editing efficiency

To assess prime editing efficiencies at the *EMX1* and *FANCF* loci, we cloned paired pegRNAs/ngRNAs into individual vectors. For lentivirus co-infection testing, we first infected MCF7 cells with EF1a-nCas9/RT lentivirus followed by treatment with puromycin (2.5 µg/ml; Sigma-Aldrich, P8833) for 2 weeks to eliminate uninfected cells. Then, EF1a-nCas9/RT-infected cells were seeded in 24-well plates at 12,500 cells per well for pegRNA and ngRNA co-infection. The infected cells were treated with hygromycin B (200 μg/ml; Gibco, 10687010) 48 hours after infection, and were collected one week after infection for editing efficiency assessment. For testing in the MCF7-nCas9/RT clonal line, we seeded cells in 24-well plates at 12,500 cells per well, followed by lentiviral infection (pegRNA-mCherry and ngRNA-EGFP). Two days after infection, mCherry and EGFP double-positive cells were isolated by FACS and cultured. Cultured cells were then collected at 2 weeks and 4 weeks post-infection for editing efficiency assessment. Genomic DNA was then extracted from each sample using the Wizard genomic DNA purification kit (Promega, A1120). Genomic sites of interest were amplified from purified genomic DNA and amplicons were sequenced on the Illumina NovaSeq 6000 platform. Briefly, sequencing libraries were prepared using DNA primers amplifying target genomic loci of interest for the first round of PCR (PCR1). Then, DNA primers containing index adapters were used for the second round of PCR (PCR2) to add these adapters to PCR1 amplicons. Finally, dual indexing primers were used for the third round PCR (PCR3) to add Illumina indexes to each PCR2 amplicon. Alignment of amplicons to reference sequences was performed using CRISPResso2^47^. For all prime editing efficiency quantification, wild-type and edited amplicon frequencies were quantified using a 21 bp window centered on either the 1 bp wild-type or edited sequence. The remaining amplicons were classified as indels.

### SNP prioritization

We selected 14 MCF7 growth-related genes overlapping with GWAS identified breast cancer susceptibility genes^26^. For each gene, we selected SNPs using the GWAS results from the Breast Cancer Association Consortium^25^. We identified genome-wide significant SNPs with GWAS *P* < 1x10^-5^, minor allele frequency < 0.02, and odds ratios < 0.9 or > 1.2 (representing approximately the top and bottom quartiles of the odds ratio distribution for SNPs meeting the location, *P* value, and MAF thresholds) for association with breast cancer within the locus +/- 500 kb of each transcription start site. We also separately selected SNPs with GWAS *P* < 1x10^-5^ in the *ESR1* locus using GWAS results from a Latina population^48^. We determined linkage disequilibrium (LD) clumps among the selected SNPs using the LD Link R package^49^ with an LD threshold of R^2^ > 0.1. We then prioritized the most likely causal variants using CAVIAR^50^, as those with a causal posterior probability (> 0.1), the highest posterior probability (≤ 0.1), or most extreme odds ratio in each haplotype block. We ran CAVIAR twice for each locus, once assuming only one causal variant per LD clump, and again allowing for more than one causal variant in each LD clump.

### Clinical variant prioritization

We retrieved clinical variants from the ClinVar database (accessed 2021-12-25), and all single nucleotide variants (SNVs) were kept for the PRIME design (**Supplementary Fig. 3c**). We first selected only the SNVs whose genes overlapped with breast cancer risk and MCF7 growth-related genes. Next, we only retained SNVs in the benign, pathogenic and uncertain significance categories. Further, for SNVs associated with *BARD1*, *BRCA1*, *BRCA2*, *RAD51C*, *RAD51D*, and *PTEN*, we only retained the SNVs with more than three submitters, as there are thousands of identified variants for these genes. Finally, our selection criteria yielded 5310 SNVs, of which we successfully designed pegRNA/ngRNA pairs for 3699 SNVs.

### Design and construction of prime-editing libraries

For nucleotide-resolution analyses of *MYC* enhancer function, paired pegRNAs/ngRNAs targeting a 716 bp enhancer region were first designed using PrimeDesign’s PooledDesign-Saturation mutagenesis tool^51^. We optimized pegRNAs/ngRNAs pairs based on ngRNA pegRNA proximity (more than 50 bp) and primer binding site (PBS) length (near 14 nt), redesigning the sequence containing the BsmBI cutting sites (GAGACG, CGTCTC) or TTTTT. Next, we used GuideScan2 to assess the specificity and efficiency of each pegRNA and ngRNA spacer sequence. Spacer sequences with low specificity were redesigned to improve the specificity. Finally, three different pegRNA/ngRNA pairs were designed to target the same base pair for 93.0% (666/716) of the substitutions. Each replicate pegRNA/ngRNA pair shared the same pegRNA and sgRNA spacer sequences, and only the substitution alleles differed in the pegRNA extension sequence. To design positive control guides, we used pegIT^52^ to generate pegRNA/ngRNA pairs which alter a single base pair to introduce a stop codon within the *MYC* coding region. We selected the best pegRNA/ngRNA pair for each position suggested by pegIT^52^. The *AAVS1* locus was selected as the targeting pegRNA/ngRNA pair negative control region based on previous work^53^, and guides were designed as described above using PrimeDesign^51^. For non-targeting pegRNA/ngRNA pairs, pegRNA and ngRNA spacer sequences and pegRNA extension sequences were selected from the ENCODE non-targeting sgRNA reference data set (https://www.encodeproject.org/files/ENCFF058BPG/). A guanine nucleotide was added to the 5’ end of all pegRNAs/ngRNAs with leading nucleotides other than G, to increase transcription efficiency from the U6 promoter. We used the following template to link these component sequences: 5’-CTTGGAGAAAAGCCTTGTTT[ngRNA-spacer]GTTTAGAGACG[5nt-random-sequence]CGTCTCACACC[pegRNA-spacer]GTTTTAGAGCTAGAAATAGCAAGTTAAAATAAGGCTAGTCCGTTATCAACTTGAAAAAGTGGCACCGAGTCGGTGC[pegRNA extension]CCTAACACCGCGGTTC-3’.

Library oligos for the *MYC* enhancer screen were synthesized by Twist Bioscience and amplified using the NEBNext High-Fidelity 2× PCR Master Mix (NEB, M0541L), forward primer: GTGTTTTGAGACTATAAATATCCCTTGGAGAAAAGCCTTGTTT and reverse primer CTAGTTGGTTTAACGCGTAACTAGATAGAACCGCGGTGTTAGG. To amplify paired PegRNA/ngRNA library oligos for enhancer saturation mutagenesis, we employed emulsion PCR (ePCR) to reduce recombination of similar amplicons during PCR. Briefly, ninety-six 20 μl ePCR reactions were performed using 0.01 fmol of pooled oligos with NEBNext High-Fidelity 2× PCR Master Mix (NEB, M0541S). Each 20 μl PCR mix was combined with 40 μl of oil-surfactant mixture (containing 4.5 % Span 80 (v/v), 0.4 % Tween 80 (v/v) and 0.05 % Triton X-100 (v/v) in mineral oil)^54^. This mixture was vortexed at maximum speed for 5 min, briefly centrifuged, and placed into the PCR machine for amplification. Thermocycler settings were: 98 °C for 30 s, then 26 cycles (98 °C 10 s, 60 °C 20 s, 72 °C 30 s), then 72 °C for 5 min, and finally a 4 °C hold. The ramp rate for each step was 2°C/s. After PCR, individual reactions were combined and purified using the QIAQuick PCR Purification Kit (Qiagen, 28104) following previously established guidelines^55^. Purified PCR products were then treated with Exonuclease I (NEB, M0568L) and purified using 1× AMPure XP beads (Beckman Coulter, A63881). The isolated ePCR products were then inserted into a BsmBI–digested lentiV2-mU6-evopreQ1 vector via Gibson assembly (NEB, E2621L). The assembled products were electroporated into Endura electrocompetent Escherichia coli cells (Biosearch Technologies, 60242) and approximately 4,000 independent bacterial colonies were cultured for each library. The resulting plasmid DNA was linearized by BsmbI digestion, gel-purified, and ligated using T4 ligase (NEB, M0202M) to a DNA fragment containing an sgRNA scaffold and the human U6 promoter. The resulting library was electroporated into Endura electrocompetent Escherichia coli cells (Biosearch Technologies, 60242) and cultured as described above. The final plasmid library was extracted using the Qiagen EndoFree Plasmid Mega Kit (Qiagen, 12381).

For the SNP and clinical variant screen Alt library, pegRNA/ngRNA pairs were designed using PrimeDesign^51^. The sequences 200 bp upstream and downstream of each variant or iSTOP were used as inputs for PrimeDesign. We generated initial pegRNA/ngRNA pairs using the following parameters: number of pegRNAs per edit: 10, length of homology downstream: 10 nt, PBS length: 13 nt, maximum reverse transcription template (RTT) length: 50 nt, number of ngRNAs per pegRNA: 10, ngRNA to pegRNA nicking distance: 50 and 75 bp. Next, a guanine nucleotide was added to the 5’ end of all pegRNAs/ngRNAs with leading nucleotides other than G to increase transcription efficiency from the U6 promoter. pegRNA/ngRNA pairs containing BsmBI sites (GAGACG, CGTCTC) or a TTTTT sequence in the pegRNA spacer, ngRNA spacer or pegRNA extension were eliminated. pegRNA/ngRNA pairs were further selected to maximize specificity, efficiency, and ngRNA to pegRNA distance while minimizing pegRNA to edit distance when multiple pairs were available for the same locus. For non-targeting pegRNA/ngRNA pairs, pegRNA spacer, ngRNA spacer and pegRNA extension sequences were selected from the ENCODE non-targeting sgRNA reference data set (https://www.encodeproject.org/files/ENCFF058BPG/). To design the Ref library, we used the same pegRNA/ngRNA pairs as the Alt library, but replaced the alternative alleles in the pegRNA extension sequences with the reference allele sequences. The final oligos adhered to the following template architecture: 5′-CTTGTGGAAAGGACGAAACACC[ngRNA-spacer]GTTTCGAGACG[6nt-random-sequence]CGTCTCTTGTTT[pegRNA-spacer]gttttagagctagaaatagcaagttaaaataaggctagtccgttatcaacttgaaaaagtggcaccgagtcggtgc[pegR NA extension]TTGACGCGGTTCTATCTAGTTAC-3′.

The Alt and Ref library oligos were synthesized by Twist Bioscience. The Alt and Ref plasmid libraries were cloned separately using two-step cloning. First, the oligo pool for each library was amplified with NEBNext High-Fidelity 2× PCR Master Mix (NEB, M0541L) and the following primers: Forward primer: TCGATTTCTTGGCTTTATATATCTTGTGGAAAGGACGAAACAC, Reverse primer: ATTTCTAGTTGGTTTAACGCGTAACTAGATAGAACCGCGTCAA. PCR products were purified via gel excision and column purification (Promega, A9282), followed by insertion into the BsmBI–digested lentiV2-hU6-evopreQ1 vector by Gibson assembly (NEB, E2621L). The assembled products were electroporated into Endura electrocompetent Escherichia coli cells (Biosearch Technologies, 60242). About 25 million bacterial colonies were cultured for each library, followed by purification with the QIAGEN Plasmid Maxi Kit (GIAGEN, 12163). For the second step, the resulting plasmid libraries from the first cloning step were linearized by BsmbI digestion, gel-purified, and ligated using T4 ligase (NEB, M0202M) to a DNA fragment containing an sgRNA scaffold and the mouse U6 promoter. The ligated products were electroporated into Endura electrocompetent Escherichia coli cells (Biosearch Technologies, 60242), and about 40 million bacterial colonies were cultured for each library. The final plasmid libraries were extracted with the Qiagen EndoFree Plasmid Mega Kit (Qiagen, 12381).

### Lentivirus production and titration

To produce the lentiviral library, we used our previously described method^46^. Briefly, 5 μg of plasmid library, with 3 μg of psPAX (Addgene, 12260) and 1 μg of pMD2.G (Addgene, 12259) packaging plasmids were cotransfected into 8 million HEK293T cells in a 10-cm dish supplemented with 36 μl PolyJet (SignaGen Laboratories, SL100688). The medium was replaced 12 hours after transfection and harvested every 24 hours thereafter for a total of three harvests. Harvested viral media was filtered through a Millex-HV 0.45-μm polyvinylidene difluoride filter (Millipore, SLHV033RS) and further concentrated via centrifugation using 100,000 NMWL (nominal molecular weight limit) Ultra-15 centrifugal filter units (Amicon, UFC910008).

The lentiviral titer was determined by transducing 400,000 cells with increasing volumes (0, 1, 2, 5, 10, 20, and 40 μl) of concentrated virus and polybrene (6 μg/ml; Millipore, TR-1003-G). 48 hours after the transduction, cells were dissociated with Trypsin-EDTA (0.25%; Gibco, 25200056) and seeded as two separate replicates; one treated with hygromycin B (200 μg/ml; Gibco, 10687010) for four days, and another that was not. Finally, hygromycin-resistant and control cells were counted to calculate the infected cell ratios and viral titers.

### Prime-editing screens

We performed *MYC* enhancer screens in triplicate. We transfected MCF7-dCas9/RT cells with lentivirus libraries at a multiplicity of infection (MOI) of 0.3 with a coverage of 1,000 transduced cells per paired pegRNA/ngRNA. 48 hours later, approximately 10 million cells were harvested as controls and the remaining cells were treated with hygromycin B (200 μg/ml; Gibco, 10687010) for 7 days. After antibiotic selection, the cells were maintained in DMEM supplemented with 10% FBS for 30 days post infection, and 10 million cells were collected from the final cell population.

We performed Alt and Ref library screens in quadruplicate. We separately infected about 24 million MCF7-nCas9/RT cells with the lentivirus library for each replicate of the Alt and Ref screens at an MOI of 0.5, with a cell coverage of 2,000 infected cells per pegRNA/ngRNA pair. 48 hours post infection, one-third of the infected cells were collected from each cell pool as control samples (Day 2). The remaining cells were treated with hygromycin B (200 μg/ml; Gibco, 10687010) for 7 days and cultured until 32 days post infection (Day 32).

### Generation of Illumina sequencing libraries

Genomic DNA was extracted from each sample via cell lysis and digestion [100 mM tris-HCl (pH 8.5), 5 mM EDTA, 200 mM NaCl, 0.2% SDS, and proteinase K (100 μg/ml)], phenol:chloroform (Thermo Fisher Scientific, 17908) extraction, and isopropanol (Thermo Fisher Scientific, BP2618500) precipitation. For the *MYC* enhancer screen, we applied ePCR during library preparation to amplify the paired pegRNA/ngRNA sequences from each sample and reduce recombination between similar sequences. Briefly, thirty 20 μl ePCRs were performed using 400 ng of DNA for each reaction and NEBNext High-Fidelity 2× PCR Master Mix (NEB, M0541S) with the following primers: Enh-lib-Forward: TCCCTACACGACGCTCTTCCGATCTNNNNNCCTTGGAGAAAAGCCTTGTTT, Enh-lib-Reverse: GGAGTTCAGACGTGTGCTCTTCCGATCTNNNNNGAACCGCGGTGTTAGG. Epcr was performed as described previously to amplify pegRNA/ngRNA pairs from genomic DNA. Thermocycler settings were 98 °C for 30 s, then 25 cycles (98 °C 10 s, 60 °C 20 s, 72 °C 1 min), then 72 °C 5 min, and finally a 4 °C hold. The ramp rate for each step was 2°C/s. After PCR, individual reactions were combined and purified using the QIAQuick PCR Purification Kit (Qiagen 28104) following previously established guidelines^55^. Purified PCR products were then treated with Exonuclease I (NEB, M0568L) and purified using 1× AMPure XP beads (Beckman Coulter, A63881). Round one PCR amplicons were used in the 2nd round of PCR to add Illumina adapter and index sequences. For the 2nd round PCR, we performed 6 ePCR reactions containing 0.023 ng of purified DNA each, using NEBNext High-Fidelity 2× PCR Master Mix (NEB, M0541S). The 2nd round PCR mixture was prepared and purified similarly to the 1st. Thermocycler settings were 98 °C for 30 s, then 12 cycles (98 °C 10 s, 60 °C 20 s, 72 °C 1 min), then 72 °C 5 min, and finally a 4 °C hold. The ramp rate for each step was 2°C/s. For Alt and Ref screens, we amplified pegRNA/ngRNA pair sequences from each sample using NEBNext High-Fidelity 2× PCR Master Mix (NEB, M0541L) and the following primers: Alt-Ref-lib-Forward: TCCCTACACGACGCTCTTCCGATCTNNNNNCTTGTGGAAAGGACGAAACACC, Alt-Ref-lib-Reverse: GGAGTTCAGACGTGTGCTCTTCCGATCTNNNNNCGTAACTAGATAGAACCGCGTCAA. Twenty-four 50 μl PCR reactions, each containing 600 ng genomic DNA, were performed for each sample. Individual reactions were combined for each sample and column purified (Promega, A9282). The purified products were then amplified by indexing PCR to add Illumina TruSeq adaptors and sample index sequences with the following primers: Index-Forward: aatgatacggcgaccaccgagatctacac[8 bp index]acactctttccctacacgacgctcttccgatct, Index-Reverse: caagcagaagacggcatacgagat[8 bp index]gtgactggagttcagacgtgtgctcttccgatct. The final libraries were gel purified and sequenced with 150 bp paired-ends on the Illumina NovaSeq 6000 platform.

### Data processing and analysis of prime-editing data

Sequencing libraries were first trimmed with 5 bp random sequences from read1 and read2, and low quality reads were filtered out with the fastp tool before formal mapping. To calculate the read counts, each pegRNA/ngRNA pair was included if it met the following criteria: (1) Read 1 exactly matched the sequence containing a 20-21 nt ngRNA spacer and 5 bp flanking sequences; (2) Read 2 exactly matched the reverse complementary sequence containing the full pegRNA extension and 5 bp flanking sequences.

For PRIME of *MYC* enhancer, the MAGeCK (0.5.9) pipeline^13^ was used to estimate the statistical significance and fold change for each pegRNA/ngRNA pair at the sgRNA level, and for each substitution at the gene level in the cell population relative to controls. The non-targeting and AAVS1 targeting pegRNAs were used as negative controls for normalization. To identify the core enhancer region for the *MYC* enhancer based on the screening results, we first identified base pairs with three significant substitutions (FDR < 0.05), and calculated the slopes for each continuous bin (moving step = 1 bp, bin size = 30 bp, x axis: the position of each base pair, y axis: the accumulation number of SBPs with three significant substitutions) (**Supplementary Fig. 2e**). The slopes were then transformed into Z score-derived *P* values accordingly. The core enhancer region was identified by merging overlapping significant bins (*P* value < 0.05).

For Alt and Ref library screens, oligos with zero reads for any sample were removed before the following analysis. Oligo counts from all samples were passed into DESeq2 (1.38.0)^31^ and a median-of-ratios method was used to normalize samples for varying sequencing depths. Normalized read counts for each oligo were then modeled by DESeq2 as a negative binomial distribution. We then used DESeq2 to check the fold changes for each oligo in Alt and Ref libraries by comparing Day 32 to Day 2 data (design= ∼ Replicate + Condition). We further estimated relative effects between the reference and alternate alleles by adding an interaction term (design=∼ Replicate + Condition + Allele + Condition:Allele). Condition refers to the collection timepoint (i.e. Day 32 or Day 2), and Allele refers to the allele category (i.e. Alt or Ref). Finally, a Wald test was performed via DESeq2 to calculate the *P* value. To minimize false positive hits and achieve an empirical FDR less than 5%, we then selected a *P* value cutoff corresponding to the fifth percentile of *P* values from non-targeting control oligos.

### Motif matrix comparison analysis

To identify potential transcription factor (TF) binding sites within the target *MYC* enhancer, we established a new method based on motif comparison^56^ to directly compare known TF motifs with our base-pair resolution functional data. We first calculated the log_2_(fold change) for each substitution at each base pair with MAGeCK (0.5.9)^13^. The log_2_(fold changes) of the wild type alleles were set to 0. We then transformed the log_2_(fold change) of each substitution into the corresponding fold change value. We further constructed the position weight matrix by normalizing the fold change of each allele per base pair to the sum of all unique alleles’ fold change per base pair. We further partitioned the enhancer sequence into multiple bins with lengths of 5 and 10 base pairs. We only retained bins with an information content (IC) over 3 and an ‘N’ content less than 10%. We then collected all TF motifs from JASPAR, HOCOMOCO, and SwissRegulon databases with high expression in MCF7 cells (TPM > 10, GSE175204). Next, we compared the filtered TF motif matrices with the enhancer bin matrix using Tomtom (*P* value < 0.05) to identify the potential TF binding sites at the enhancer. Finally, we only retained positive TF motif hits overlapping at least 95% of the input sequences’ essential base pairs (positions with maximum probabilities > 0.5). Details about the best matching motifs are summarized in **Supplementary Table 2**.

### Predicting base pair contribution to enhancer activity with BPNet

We trained a convolutional neural network using BPNet consistent with the published approach^24^ to explain the GATA3, ELF1, FOXM1, MTA3, and RCOR1 ChIP-seq data from ENCODE projects. Briefly, the model inputs were 1kb sequences across each ChIP-seq peak locus, and corresponding ChIP-seq control peaks were used as the bias track for training. The region from chromosome 2 was used as the tuning set, and chromosomes 5, 6, 7, 10, and 14 were used as the test set. The X and Y chromosomes were excluded. The remaining regions from other chromosomes were used to train the model with default parameters. Once models were acquired for each TF’s ChIP-seq data, DeepLIFT was used to calculate each input sequence base pair’s contribution to enhancer activity. TF-MoDISco contribution scores were finally used to cluster and determine consolidated TF motifs and map these to input peak regions.

### MCF7 genotyping analysis

Sequence Read Archive (SRA) files for SRR7707725 and SRR7707726 (paired-end, two reads per loci) were retrieved from BioProject PRJNA486532. We used bwa-mem v.0.7.17 to align sequenced reads to the human reference genome hg38 for each run separately. The Picard tools, SortSam, MarkDuplicates, AddOrReplaceReadGroups were then used to process the BAM files. Finally, GATK v.4.2.5.0 was used to call SNPs and indels via local haplotype re-assembly (HaplotypeCaller) followed by joint genotyping on a single-sample GVCF from HaplotypeCaller (GenotypeGVCFs). Finally, CalcMatch v.1.1.2 was used to verify genotype consistency between two runs.

### Motif scan and TF identification for alleles with functional breast cancer SNPs

The sequences 20 bp upstream and downstream of each SNP (Alt and Ref alleles) were used as input sequences for TF motif analysis. FIMO software (version 5.5.0)^57^ was used to identify matching motifs centered on the SNP regions against the human TF motif database HOCOMOCO (v11 FULL)^19^. All FIMO motif scans were performed using default settings. Finally, TFs (FPKM >1) with binding motifs overlapping target SNP loci were selected (FDR < 0.05, *P* value < 0.0001).

### Functional validation of rs10956415 using prime editing and RT-qPCR

To validate the function of rs10956415 in MCF7 cells, we converted the alternative allele (A) to the reference allele (C) at this locus using PE. To clone the ngRNA/pegRNA expression plasmid, we amplified the fragment containing the ngRNA-mU6-pegRNA for the rs10956415 reference allele (C) from the screening plasmid library, and inserted this fragment into the BsmBI–digested lentiV2-hU6-evopreQ1 vector using Gibson assembly (NEB, E2621L). We verified the cloned ngRNA/pegRNA plasmid sequence using Primordium whole-plasmid sequencing.

To perform PE, we transfected two million MCF7-dCas9/RT cells with 2000 ng of ngRNA/pegRNA plasmid containing an EGFP marker using PolyJet (SignaGen Laboratories, SL100688). Five days after transfection, we sorted the cells with the highest EGFP expression level (top 2%) into 96-well plates with 100 cells per well using FACS. Approximately two weeks later, we extracted genomic DNA from half of the cells in each well and maintained the other half by seeding them in a 24-well plate. We estimated the PE efficiency for each well by performing genotyping PCR followed by Sanger sequencing. We then expanded the cells in the wells with the highest editing efficiency to isolate clonal PE edited cell lines. We sorted the cell pool into 96-well plates with one cell per well using FACS. Approximately two weeks later, we performed genotyping PCR followed by Sanger sequencing to identify successfully edited clones. Deep sequencing was then performed to quantify the copy number of edited alleles.

To assess the effect of rs10956415 on *MYC* expression, we used seven PE edited clones with four copies of the C allele and one copy of the A allele. About two million cells from each sample were used to extract total RNA with the RNeasy Plus Mini Kit (Qiagen, #74134), and 1 µg of RNA was used to generate cDNA with the iScript cDNA Synthesis Kit (Bio-Rad, #1708890). We used RT-qPCR to quantify *MYC* expression (forward primer: GTCCTCGGATTCTCTGCTCT, reverse primer: ATCTTCTTGTTCCTCCTCAGAGTC), which was normalized to the *GAPDH* expression level (forward primer: CCACTCCTCCACCTTTGACG, reverse primer: ATGAGGTCCACCACCCTGTT).

### Protein structure prediction with AlphaFold

To explore the impact of the BARD1 His36Pro mutation on BARD1/BRCA1 complex structure, we predicted the wild type BRAD1/BRCA1 and BARD1(His36Pro)/BRCA1 complex structures with AlphaFold. We used the same amino acid chain which is used in the BARD1/BRCA1 complex structure determined by NMR spectroscopy^42^ (BARD1, residues 26-122; BRCA1, residues 1-103) as input for complex structure predictions. The amino acid chains of BARD1 and BRCA1 were imported into the Google Colab Version of AlphaFold V2.2.4^58, 59^, powered by Python 3 Google Compute Engine. AlphaFold applied a multimer model in response to the duo-sequence imputation, then searched the genetic database to determine the best suited multiple sequence alignment (MSA) for the imported sequence and initiated structural prediction. To avoid stereochemical violations, all structures are relaxed with AMBER model (Assisted Model Building with Energy Refinement) using GPU acceleration. The resulting PDB files were imported into UCSF Chimera X^60, 61^ for structure visualization. Protein chains were assigned different colors to distinguish individual chains, and selected amino acid atomic structures and hydrogen bonds were illustrated for interaction analysis. Finally, the real-time rendered complex structures were exported using the snapshot function in Chimera X at the optimal visualization angle.

---

**Supplementary Figure 1.**
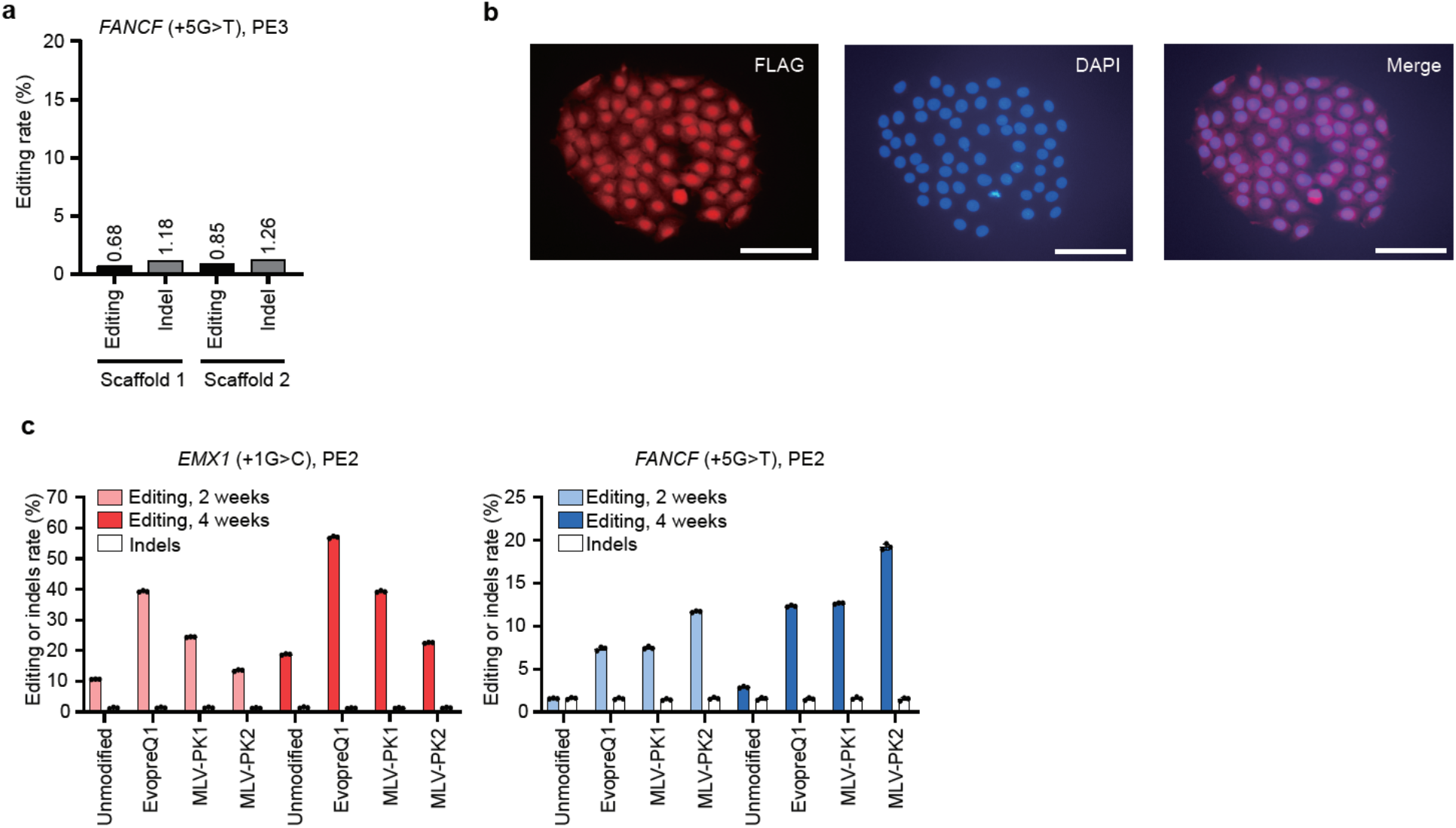
Optimizing PE efficiency in MCF7 cell line. (a) Prime editing efficiency and indel rate by co-infection of pegRNA, ngRNA and nCas9/RT expressing lentiviruses in MCF7 cells. (b) Immunofluorescent staining showing the localization of nCas9/RT (red, FLAG tagged) in the nucleus (blue, DAPI) in MCF7-nCas9/RT cells. Scale bars, 1000 μm. (c) Editing efficiency and indel rate by PE using three different structured RNA motifs to the 3’ terminus of pegRNAs at 2 and 4 weeks post infection in MCF7-nCas9/RT cells. Error bars represent the s.d.

**Supplementary Figure 2.**
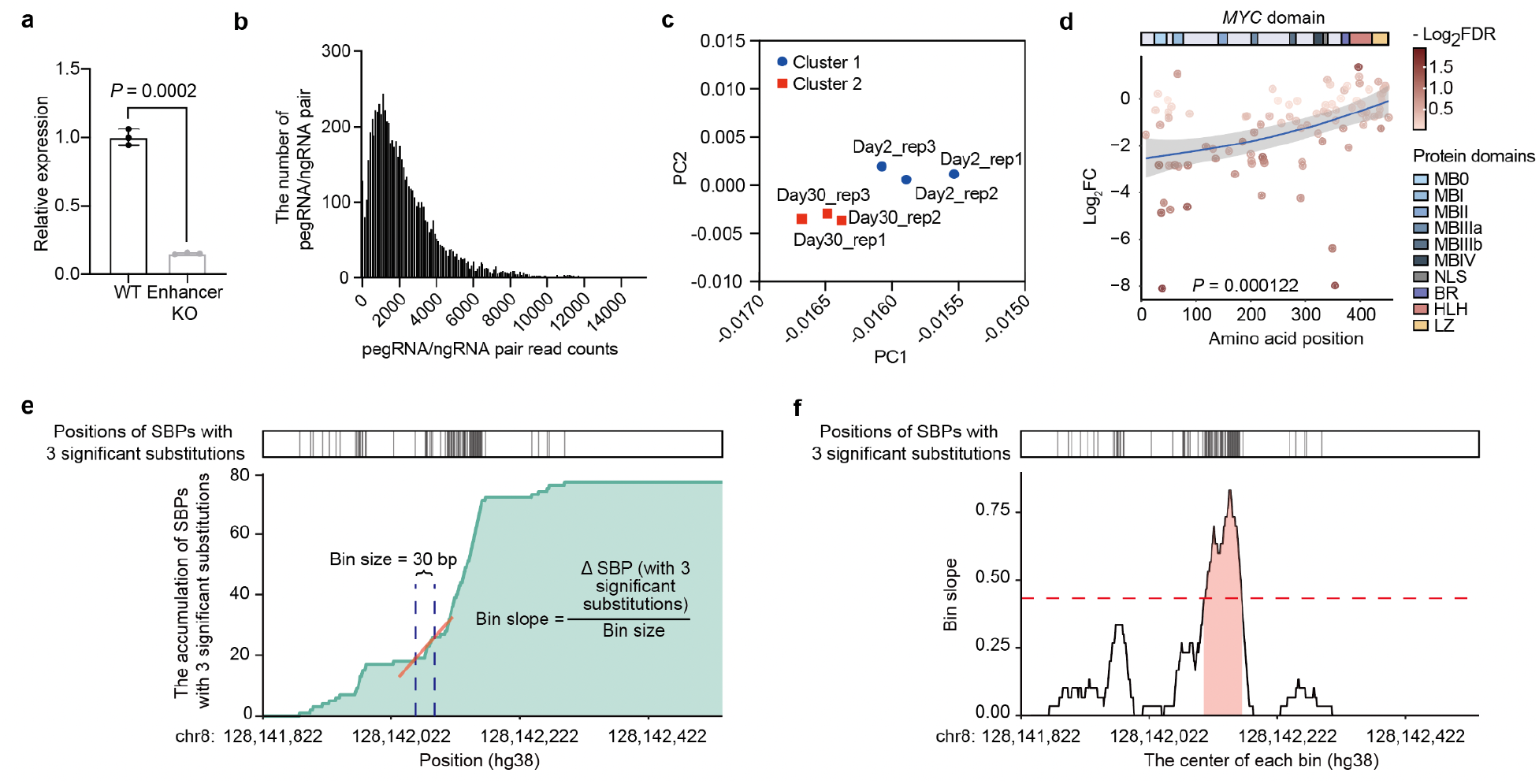
Characterize enhancer function and results of PRIME in MCF7 cells. (a) CRISPR/Cas9 knockout of the *MYC* enhancer in MCF7 decreased *MYC* expression. *P* values were calculated using a two-tailed two-sample t-test. Error bars represent the s.e.m. (b) Distribution of pegRNA/ngRNA pair read counts in the cloned plasmid library. (c) PCA analysis demonstrates the high reproducibility of PRIME between biological replicates. (d) The correlation between locations of PE-induced stop codons and their effect sizes. The blue line and *P* value were calculated using generalized additive models. The shaded areas indicate 95% confidence intervals. (e) (Top) The position of SBPs with three significant substitutions. (Bottom) Cumulative distribution plot of SBPs with three significant substitutions along the *MYC* enhancer and the formula for calculating the slope of each continuous bin. (f) Line plot of slopes for each continuous bin along the *MYC* enhancer. The red dashed line is the cutoff for a significant slope, which is based on a slope with a Z score-derived *P* value equal to 0.05. The red region is the core enhancer region, derived from the bins’ slopes greater than the cutoff (slope > 0.43).

**Supplementary Figure 3.**
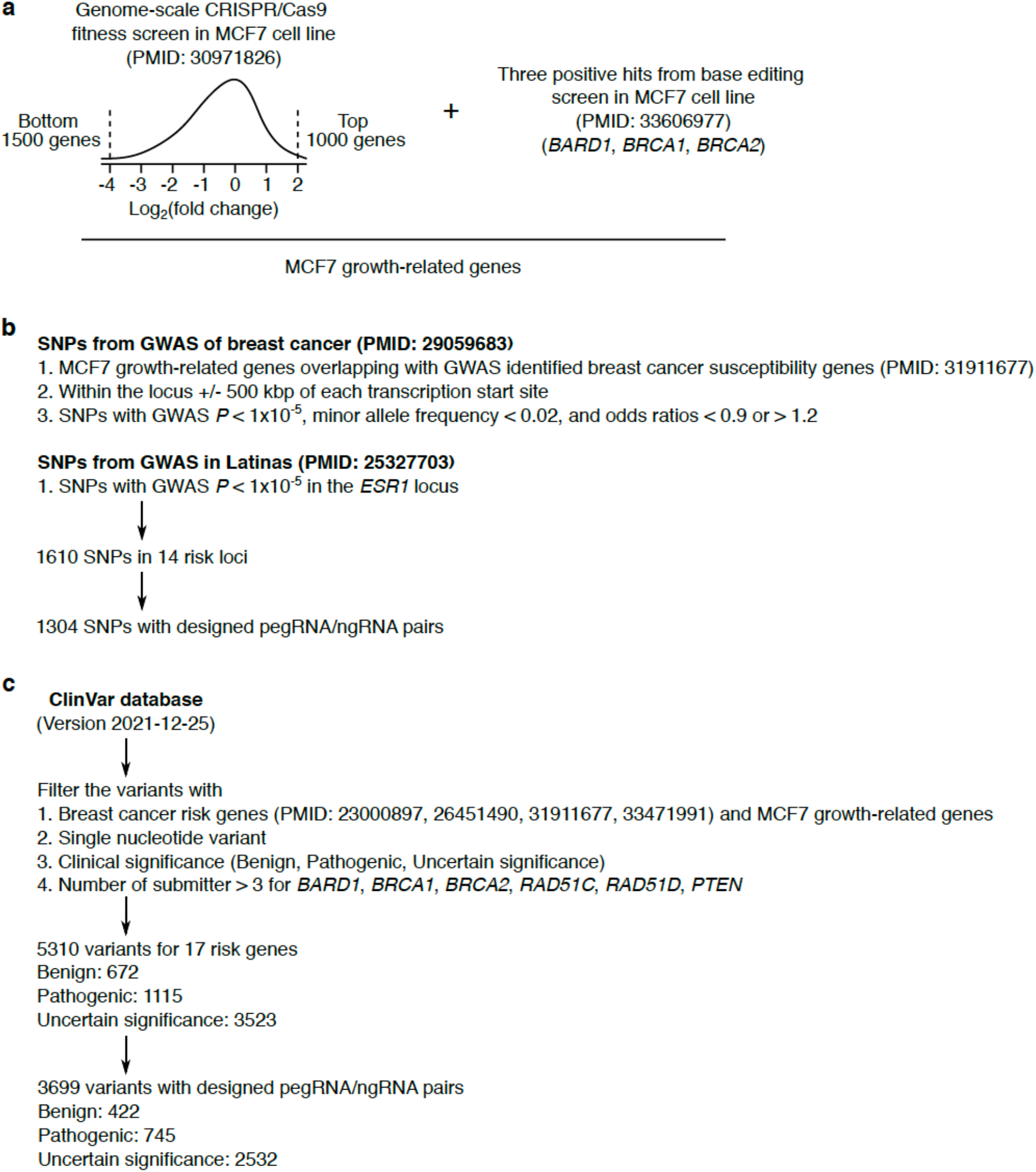
Strategies for prioritizing genomic loci and clinical variants. (a) The MCF7 growth-related genes were selected from the CRISPR/Cas9 knockout screen and base editing screen in MCF7 cells. (b) The strategy used for selecting breast cancer-related SNPs. (c) The strategy used for selecting clinical variants.

**Supplementary Figure 4.**
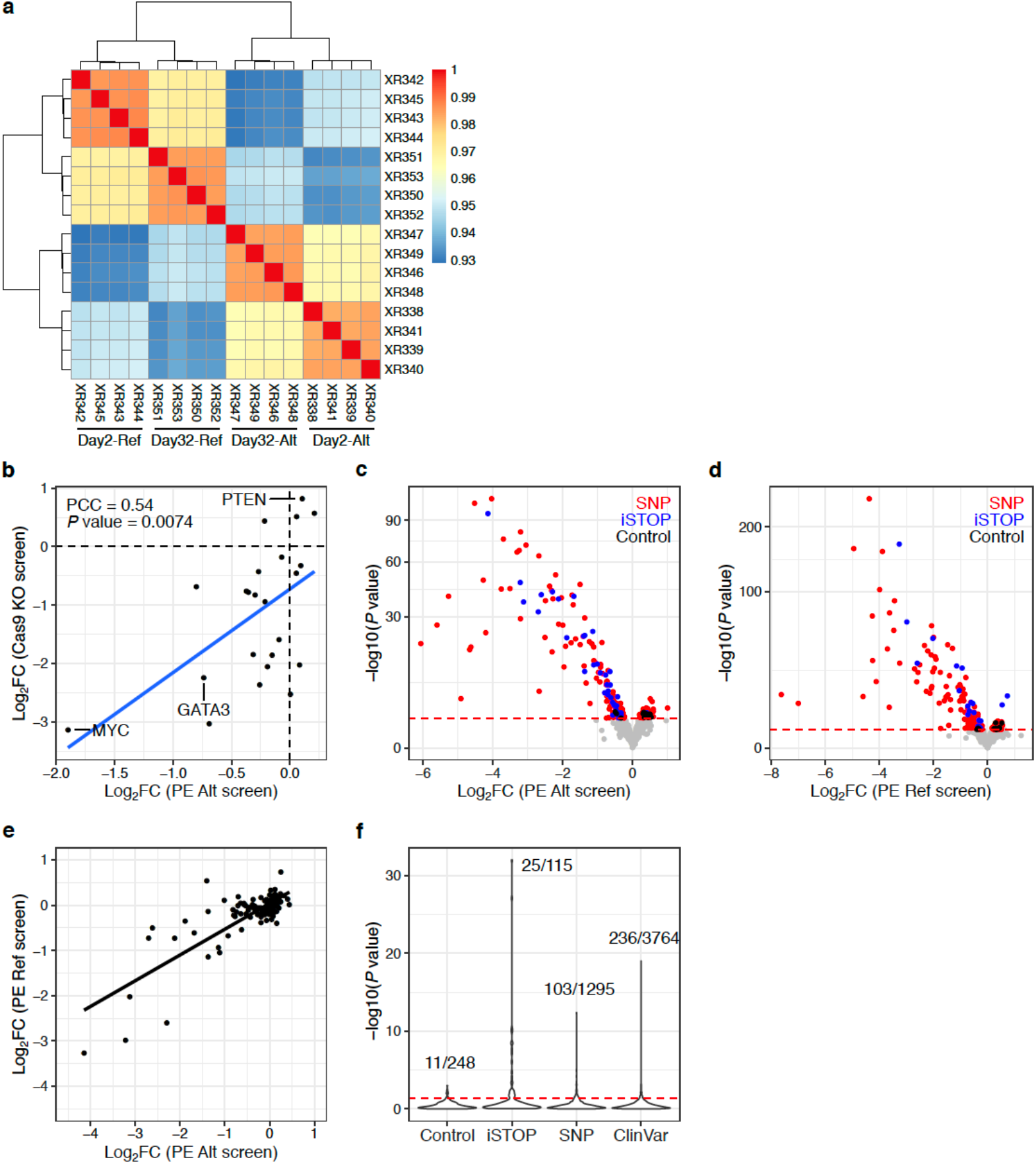
Quality control and primary analysis of disease variants. (a) Heatmap with pairwise correlations and hierarchical clustering of read counts from PRIME. (b) Pearson correlations between the log2(fold change) of iSTOPs in the Alt library screen and the log2(fold change) of gRNAs in the CRISPR/Cas9 knockout screen for each target gene. (c) Volcano plot of the results from the Alt library screen. (d) Volcano plot of the results from the Ref library screen. (e) The log2(fold change) for each iSTOP from the Alt and Ref library screens. (f) Violin plot showing the 5% FDR cutoff used for the relative effect analysis comparing the Alt and Ref libraries. Numbers above peaks indicate the significant data points versus the total data points in each category when using 5% FDR. We used the 5% percentile of *P* values from negative controls as the empirical significance threshold to achieve a false discovery rate (FDR) of 5% indicated by the red dashed line in d-f.

**Supplementary Figure 5.**
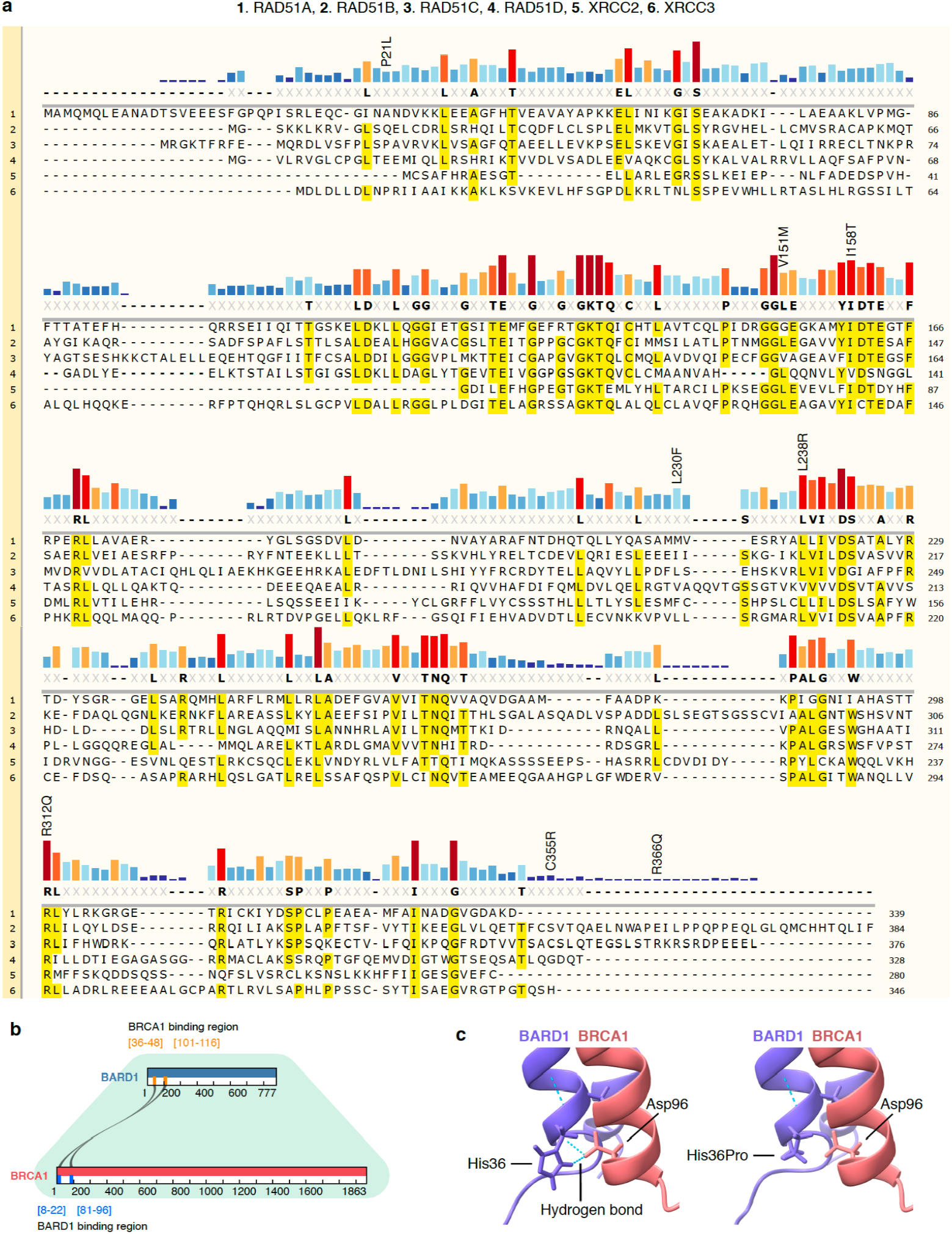
Examples of functional VUS with their potential consequences. (a) Sequence conservation of RAD51 family proteins. Alignment of RAD51 family proteins using MUSCLE. Functional VUS identified by PRIME in RAD51C are labeled. (b) Graphic showing the binding regions between BARD1 and BRCA1. (c) The Alphafold predicted protein structure of the BARD1 and BRCA1 complex. Two hydrogen bonds were identified between wild type His36 in BARD1 and Asp96 in BRCA1, but lost following the BARD1 His36Pro mutation.

## References

1. Taliun, D. et al. Sequencing of 53,831 diverse genomes from the NHLBI TOPMed Program. Nature 590, 290–299 (2021).

2. Shalem, O., Sanjana, N.E. & Zhang, F. High-throughput functional genomics using CRISPR-Cas9. Nat Rev Genet 16, 299–311 (2015).

3. Anzalone, A.V., Koblan, L.W. & Liu, D.R. Genome editing with CRISPR-Cas nucleases, base editors, transposases and prime editors. Nat Biotechnol 38, 824-844 (2020).

4. Chen, P.J. & Liu, D.R. Prime editing for precise and highly versatile genome manipulation. Nat Rev Genet (2022).

5. Anzalone, A.V. et al. Search-and-replace genome editing without double-strand breaks or donor DNA. Nature 576, 149–157 (2019).

6. Erwood, S. et al. Saturation variant interpretation using CRISPR prime editing. Nat Biotechnol 40, 885–895 (2022).

7. Anzalone, A.V., Lin, A.J., Zairis, S., Rabadan, R. & Cornish, V.W. Reprogramming eukaryotic translation with ligand-responsive synthetic RNA switches. Nat Methods 13, 453–458 (2016).

8. Houck-Loomis, B. et al. An equilibrium-dependent retroviral mRNA switch regulates translational recoding. Nature 480, 561–564 (2011).

9. Nelson, J.W. et al. Engineered pegRNAs improve prime editing efficiency. Nat Biotechnol 40, 402–410 (2022).

10. Dang, Y. et al. Optimizing sgRNA structure to improve CRISPR-Cas9 knockout efficiency. Genome Biol 16, 280 (2015).

11. Chen, P.B. et al. Systematic discovery and functional dissection of enhancers needed for cancer cell fitness and proliferation. Cell Rep 41, 111630 (2022).

12. Cho, S.W. et al. Promoter of lncRNA Gene PVT1 Is a Tumor-Suppressor DNA Boundary Element. Cell 173, 1398–1412 e1322 (2018).

13. Li, W. et al. MAGeCK enables robust identification of essential genes from genome-scale CRISPR/Cas9 knockout screens. Genome Biol 15, 554 (2014).

14. Shalem, O. et al. Genome-scale CRISPR-Cas9 knockout screening in human cells. Science 343, 84–87 (2014).

15. Baluapuri, A., Wolf, E. & Eilers, M. Target gene-independent functions of MYC oncoproteins. Nat Rev Mol Cell Biol 21, 255–267 (2020).

16. Vitsios, D., Dhindsa, R.S., Middleton, L., Gussow, A.B. & Petrovski, S. Prioritizing non-coding regions based on human genomic constraint and sequence context with deep learning. Nat Commun 12, 1504 (2021).

17. Villar, D. et al. Enhancer evolution across 20 mammalian species. Cell 160, 554–566 (2015).

18. Fornes, O., et al. JASPAR 2020: update of the open-access database of transcription factor binding profiles. Nucleic Acids Res 48, D87–D92 (2020).

19. Kulakovskiy, I.V. et al. HOCOMOCO: towards a complete collection of transcription factor binding models for human and mouse via large-scale ChIP-Seq analysis. Nucleic Acids Res 46, D252–D259 (2018).

20. Pachkov, M., Balwierz, P.J., Arnold, P., Ozonov, E. & van Nimwegen, E. SwissRegulon, a database of genome-wide annotations of regulatory sites: recent updates. Nucleic Acids Res 41, D214–220 (2013).

21. Consortium, E.P. An integrated encyclopedia of DNA elements in the human genome. Nature 489, 57–74 (2012).

22. Schreiber, J., Durham, T., Bilmes, J. & Noble, W.S. Avocado: a multi-scale deep tensor factorization method learns a latent representation of the human epigenome. Genome Biol 21, 81 (2020).

23. Behan, F.M. et al. Prioritization of cancer therapeutic targets using CRISPR-Cas9 screens. Nature 568, 511–516 (2019).

24. Avsec, Z. et al. Base-resolution models of transcription-factor binding reveal soft motif syntax. Nat Genet 53, 354–366 (2021).

25. Michailidou, K. et al. Association analysis identifies 65 new breast cancer risk loci. Nature 551, 92–94 (2017).

26. Fachal, L. et al. Fine-mapping of 150 breast cancer risk regions identifies 191 likely target genes. Nat Genet 52, 56–73 (2020).

27. Hanna, R.E., et al. Massively parallel assessment of human variants with base editor screens. Cell 184, 1064-1080 e1020 (2021).

28. Landrum, M.J. et al. ClinVar: improvements to accessing data. Nucleic Acids Res 48, D835–D844 (2020).

29. Cuella-Martin, R. et al. Functional interrogation of DNA damage response variants with base editing screens. Cell 184, 1081–1097 e1019 (2021).

30. Qi, L.S. et al. Repurposing CRISPR as an RNA-guided platform for sequence-specific control of gene expression. Cell 152, 1173–1183 (2013).

31. Love, M.I., Huber, W. & Anders, S. Moderated estimation of fold change and dispersion for RNA-seq data with DESeq2. Genome Biol 15, 550 (2014).

32. Bruna, A. et al. TGFbeta induces the formation of tumour-initiating cells in claudinlow breast cancer. Nat Commun 3, 1055 (2012).

33. Bossone, S.A., Asselin, C., Patel, A.J. & Marcu, K.B. MAZ, a zinc finger protein, binds to c-MYC and C2 gene sequences regulating transcriptional initiation and termination. Proc Natl Acad Sci U S A 89, 7452–7456 (1992).

34. Wang, X. et al. MAZ drives tumor-specific expression of PPAR gamma 1 in breast cancer cells. Breast Cancer Res Treat 111, 103–111 (2008).

35. Ghandi, M. et al. Next-generation characterization of the Cancer Cell Line Encyclopedia. Nature 569, 503–508 (2019).

36. Wang, Y.H. et al. Knockdown of c-Myc expression by RNAi inhibits MCF-7 breast tumor cells growth in vitro and in vivo. Breast Cancer Res 7, R220–228 (2005).

37. Kircher, M. et al. A general framework for estimating the relative pathogenicity of human genetic variants. Nat Genet 46, 310–315 (2014).

38. Pollard, K.S., Hubisz, M.J., Rosenbloom, K.R. & Siepel, A. Detection of nonneutral substitution rates on mammalian phylogenies. Genome Res 20, 110–121 (2010).

39. Li, W. et al. A synergetic effect of BARD1 mutations on tumorigenesis. Nat Commun 12, 1243 (2021).

40. UniProt, C. UniProt: the universal protein knowledgebase in 2021. Nucleic Acids Res 49, D480–D489 (2021).

41. Prakash, R. et al. Homologous recombination-deficient mutation cluster in tumor suppressor RAD51C identified by comprehensive analysis of cancer variants. Proc Natl Acad Sci U S A 119, e2202727119 (2022).

42. Brzovic, P.S., Rajagopal, P., Hoyt, D.W., King, M.C. & Klevit, R.E. Structure of a BRCA1-BARD1 heterodimeric RING-RING complex. Nat Struct Biol 8, 833–837 (2001).

43. Densham, R.M. et al. Human BRCA1-BARD1 ubiquitin ligase activity counteracts chromatin barriers to DNA resection. Nat Struct Mol Biol 23, 647–655 (2016).

44. Spain, B.H., Larson, C.J., Shihabuddin, L.S., Gage, F.H. & Verma, I.M. Truncated BRCA2 is cytoplasmic: implications for cancer-linked mutations. Proc Natl Acad Sci U S A 96, 13920–13925 (1999).

45. Mandegar, M.A. et al. CRISPR Interference Efficiently Induces Specific and Reversible Gene Silencing in Human iPSCs. Cell Stem Cell 18, 541–553 (2016).

46. Ren, X. et al. Parallel characterization of cis-regulatory elements for multiple genes using CRISPRpath. Sci Adv 7, eabi4360 (2021).

47. Clement, K. et al. CRISPResso2 provides accurate and rapid genome editing sequence analysis. Nat Biotechnol 37, 224–226 (2019).

48. Fejerman, L. et al. Genome-wide association study of breast cancer in Latinas identifies novel protective variants on 6q25. Nat Commun 5, 5260 (2014).

49. Machiela, M.J. & Chanock, S.J. LDlink: a web-based application for exploring population-specific haplotype structure and linking correlated alleles of possible functional variants. Bioinformatics 31, 3555–3557 (2015).

50. Hormozdiari, F., Kostem, E., Kang, E.Y., Pasaniuc, B. & Eskin, E. Identifying causal variants at loci with multiple signals of association. Genetics 198, 497–508 (2014).

51. Hsu, J.Y. et al. PrimeDesign software for rapid and simplified design of prime editing guide RNAs. Nat Commun 12, 1034 (2021).

52. Anderson, M.V., Haldrup, J., Thomsen, E.A., Wolff, J.H. & Mikkelsen, J.G. pegIT - a web-based design tool for prime editing. Nucleic Acids Res 49, W505–W509 (2021).

53. Chen, C.H. et al. Improved design and analysis of CRISPR knockout screens. Bioinformatics 34, 4095–4101 (2018).

54. Williams, R. et al. Amplification of complex gene libraries by emulsion PCR. Nat Methods 3, 545–550 (2006).

55. Verma, V., Gupta, A. & Chaudhary, V.K. Emulsion PCR made easy. Biotechniques 69, 421–426 (2020).

56. Gupta, S., Stamatoyannopoulos, J.A., Bailey, T.L. & Noble, W.S. Quantifying similarity between motifs. Genome Biol 8, R24 (2007).

57. Grant, C.E., Bailey, T.L. & Noble, W.S. FIMO: scanning for occurrences of a given motif. Bioinformatics 27, 1017–1018 (2011).

58. Jumper, J. et al. Highly accurate protein structure prediction with AlphaFold. Nature 596, 583–589 (2021).

59. Mirdita, M. et al. ColabFold: making protein folding accessible to all. Nat Methods 19, 679–682 (2022).

60. Goddard, T.D. et al. UCSF ChimeraX: Meeting modern challenges in visualization and analysis. Protein Sci 27, 14–25 (2018).

61. Pettersen, E.F. et al. UCSF ChimeraX: Structure visualization for researchers, educators, and developers. Protein Sci 30, 70–82 (2021).

